# Histone demethylase complexes KDM3A and KDM3B cooperate with OCT4/SOX2 to construct pluripotency gene regulatory network

**DOI:** 10.1101/2020.08.16.245639

**Authors:** Zhenshuo Zhu, Xiaolong Wu, Qun Li, Juqing Zhang, Shuai Yu, Qiaoyan Shen, Zhe Zhou, Qin Pan, Wei Yue, Dezhe Qin, Ying Zhang, Wenxu Zhao, Rui Zhang, Sha Peng, Na Li, Shiqiang Zhang, Anmin Lei, Yi-Liang Miao, Zhonghua Liu, Xingqi Chen, Huayan Wang, Mingzhi Liao, Jinlian Hua

**Affiliations:** College of Veterinary Medicine, Shaanxi Centre of Stem Cells Engineering & Technology, Northwest A&F University, Yangling, Shaanxi, 712100, China; College of Life Science, Northwest A&F University, Yangling, Shaanxi, 712100, China; Institute of Stem Cell and Regenerative Biology, College of Animal Science and Veterinary Medicine, Huazhong Agricultural University, Wuhan, 430070, China; Key Laboratory of Animal Cellular and Genetic Engineering of Heilongjiang Province, College of Life Science, North-East Agricultural University, Harbin, China; Department of Cell and Molecular Biology, Karolinska Institutet, 17177, Solna, Sweden

**Keywords:** Pluripotency, H3K9 methylation, Histone demethylase complexes, KDM3A/3B

## Abstract

The pluripotency gene regulatory network of porcine-induced pluripotent stem cells (piPSCs), especially in epigenetics, remains elusive. To determine this biological function of epigenetics, we cultured piPSCs in different culture conditions. We found that activation of pluripotent gene- and pluripotency-related pathways requires the erasure of H3K9 methylation modification which was further influenced by mouse embryonic fibroblast (MEF) served feeder. By dissecting the dynamic change of H3K9 methylation during loss of pluripotency, we demonstrated that the H3K9 demethylases KDM3A and KDM3B regulated global H3K9me2/me3 level and that their co-depletion led to the collapse of the pluripotency gene regulatory network. Immunoprecipitation-mass spectrometry (IP-MS) provided evidence that KDM3A and KDM3B formed a complex to perform H3K9 demethylation. The genome-wide regulation analysis revealed that OCT4 (O) and SOX2 (S), the core pluripotency transcriptional activators, maintained the pluripotent state of piPSCs depending on the H3K9 hypomethylation. Further investigation revealed that O/S cooperating with histone demethylase complex containing KDM3A and KDM3B promoted pluripotency genes expression to maintain the pluripotent state of piPSCs. Together, these data offer a unique insight into the epigenetic pluripotency network of piPSCs.

**Summary:** Erasure of H3K9 methylation in porcine pluripotent stem cells depends on the complex of transcription factors OCT4/SOX2 and histone demethylase KDM3A/KDM3B.

## Background

The acquisition of pluripotency depends on the reconstruction of the epigenetic modification during reprogramming. Recent studies have shown that OCT4, SOX2, and KLF4 (OSK) cooperating with epigenetic modifiers mediated the gradually silencing of differentiation genes and activating of pluripotency genes simultaneously during reprogramming [1, 2]. Of note, global H3K9 methylation is erased in the later stages of the reprogramming with the activation of pluripotency network [3, 4]. Therefore, the establishment of pluripotent epigenetic regulation network based on histone modification should be the kernel for maintaining the pluripotent state of induced pluripotent stem cells (iPSCs) [4].

As an epigenetic modification associated with transcriptionally repressing, H3K9 methylation is a major hindrance during somatic cell reprogramming into iPSCs [3, 4]. Loss of H3K9 methylation significantly increases the expression of endogenous pluripotent genes and contributes to the transition of pre-iPSCs into iPSCs [3, 5]. Previous studies also demonstrated that the increase of H3K9 methylation causes the loss of pluripotency in mouse embryonic stem cells (mESCs) [6-8]. However, the molecular mechanism underlying H3K9 hypomethylation in pluripotent gene regions remains unclear in pluripotent stem cells. The KDM3 family, which includes KDM3A and KDM3B, function as epigenetic activators to demethylate intrinsic H3K9 methylation modification [9, 10]. Additionally, KDM3A has been found to play a critical role in a wide range of biological process like embryo development [8], pluripotency maintenance [6], carcinogenesis [11, 12], cell senescence [13], sex differentiation [14], and spermatogenesis [15, 16]. At the same time, KDM3B exhibits similar functions like KDM3A, such as pluripotency maintenance [5, 7], carcinogenesis [17], and spermatogenesis [18]. However, the elaborate mechanism of KDM3A and KDM3B in pluripotency maintenance and their functional redundancy in H3K9 demethylation remain unclear.

Pig is an ideal model for human medical research due to its close similarity with human in genetic, anatomical, and physiological condition [19, 20]. Research on porcine PSCs contributes to promising value in the biomedical field [20], though most published porcine PSCs, including ESCs and iPSCs, cannot meet with the stringent criteria for pluripotency identity. We hypothesized the unsound pluripotency gene regulatory network of piPSCs, especially in epigenetics, would be the major obstacle for us to obtain bona fide PSCs *in vitro*. To solve this issue, the present study used piPSCs generated by tetracycline operator (TetO)-inducible system [21] to delineate the mechanism of H3K9 demethylation in piPSCs. Here, we found that KDM3A/KDM3B-meditated H3K9 demethylation plays a critical role in pluripotency maintenance of piPSCs. Moreover, the rescue experiment suggested that not all exogenous genes were required and only O/S were essential for the pluripotency maintenance of piPSCs. By performing genome-wide analysis of H3K9me2/me3 modification and O/S occupancy enrichment by ChIP-seq, as well as KDM3A and KDM3B interactome analysis by IP-MS, we thus reveal the novel pluripotency regulation pathway, in which SOX2/OCT4 recruit chromatin remodeling proteins KDM3A/KDM3B to form a transcriptional complex to drive the pluripotency gene regulatory network.

## Results

### H3K9 demethylation maintains the pluripotency of piPSCs

We applied a combination of cytokines (LIF and bFGF) and signaling inhibitors (2i: chir99021 and sb431542) to maintain piPSCs that were reprogrammed by TetO-inducible systems (TetO-FUW-OSKM and FUW-M2rtTA) [21]. The piPSCs depended on the sustained expression of exogenous genes, which were similar to other reported piPSCs [21-24]. The clonal morphology and pluripotency gradually disappeared after the piPSCs cultured in differentiation medium without doxycycline (DOX) and feeder (Fig. S1A). The expression of both exogenous and endogenous pluripotent genes (*OCT4, SOX2, LIN28A*, and *NANOG*) were down-regulated in this treatment (Fig. S1B, C). However, the expression of endogenous *KLF4* was not impeded, and *c-MYC* was significantly up-regulated (Fig. S1B). Analysis of global histone methylation showed that the level of H3K9me1/2/3 was significantly increased, whereas the level of H3K4me1/3 was decreased (Fig. S1D). The RT-qPCR analysis further showed the expression of *KDM3A/3B/3C/4B/4C*, which are H3K9 demethylase, was down-regulated significantly (Fig. S1E). These results indicated that the H3K9 hypomethylation and pluripotent factors *OCT4* and *SOX2* were strongly associated with maintaining the pluripotency of piPSCs. Additionally, the downregulation of H3K9 demethylase KDM3A/3B/ 3C/4B/4C might be caused by the loss of pluripotent factors, cytokines, and small molecules during differentiation.

To investigate the mechanism in detail, we screened a series of components in the culture medium (Fig. S1F). The results showed that the pluripotency of piPSCs mainly depends on two small molecule inhibitors, DOX, and feeder (Fig. 1A, S1F). Given that the expression of *KDM3A* was significantly down-regulated during differentiation (Fig. S1E), we further tested the expression of *KDM3A* in different treatments. The results showed that the expression of *KDM3A* was affected by feeder. Interestingly, the expression of *KDM3A* was not impeded when DOX was removed, but it was further down-regulated when DOX was removed under F-treatment (Fig. S1G). These results suggested the possible pivotal role of feeder and core pluripotent factors in reducing H3K9 methylation.

**Fig. 1.**
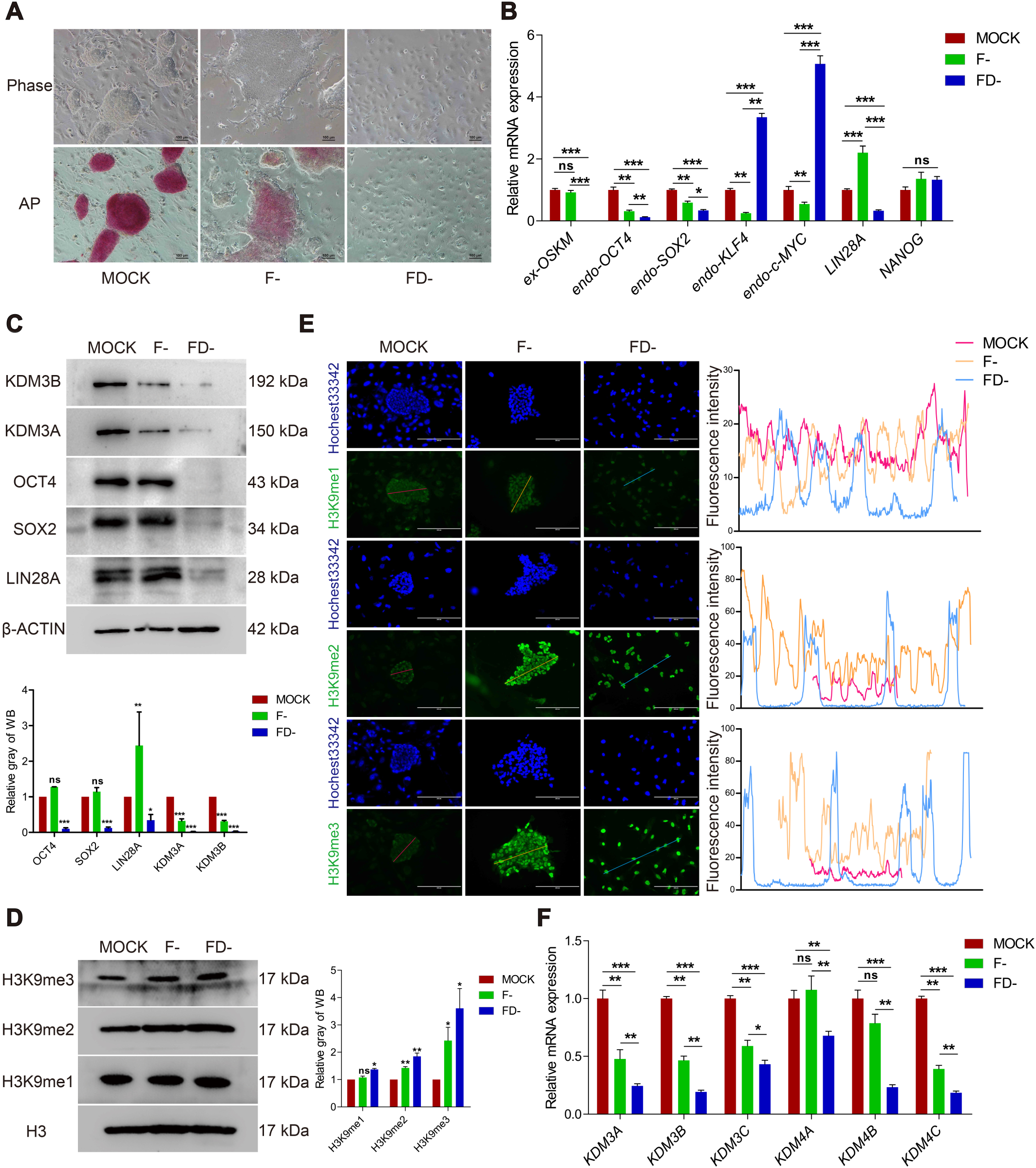
Feeder and exogenous genes affect the global H3K9 methylation in piPSCs. **(A)** Representative image of Alkaline Phosphatase (AP) stained colonies after 5 days of clonal growth in the F- (feeder free) and FD- (feeder free and absence of Dox) treatments. The MOCK represents piPSCs grown under normal condition as a blank control. The experiments were performed three times. The scale bar represents 100 μm. **(B)** RT–qPCR analysis of the exogenous reprogramming factors and endogenous pluripotent genes in the F- and FD-treatments. The relative expression levels were normalized to β-actin. Data represent the mean ± s.d.; n = 3 independent experiments. **(C)** Representative Western-Blot of OCT4, Sox2, LIN28A, KDM3A and KDM3B after 5 days of culture in the indicated conditions. The quantitative analysis is shown by histogram. Data represent the mean ± s.d.; n = 3 independent experiments. **(D)** Representative Western-Blot of H3K9me1/2/3 after 5 days of culture in the indicated conditions. The quantitative analysis is shown by histogram. Data represent the mean ± s.d.; n = 3 independent experiments. **(E)** Immunofluorescence analysis of H3K9me1/2/3 in the indicated conditions. Intensity values along the indicated path were obtained using ImageJ and shown by line chart. The nuclei were DAPI stained. The scale bar represents 200 μm. The experiments were performed three times. **(F)** RT–qPCR analysis of *KDM3A/3B/3C* and *KDM4A/4B/4C* in the F- and FD-treatments. The relative expression levels were normalized to β-actin. Data represent the mean ± s.d.; n = 3 independent experiments.

To investigate the mechanisms of increased H3K9 methylation under the feeder-free condition, we focused on the F- and FD-treatments. With the disappearance of typical clone morphology, lower proliferation rate, and high rate of apoptosis under both conditions were observed (Fig. 1A, S2A, B, C). The expression levels of *OCT4* and *SOX2* were gradually down-regulated, whereas those of *KLF4* and *c-MYC* were significantly up-regulated (Fig. 1B, C). The expression of *LIN28A* increased significantly in F-treatment but decreased significantly in FD-treatment (Fig.1B, C). Immunofluorescence also confirmed that the total expressions of OCT4 and SOX2 were significantly decreased and that the piPSCs gradually differentiated into the ectoderm and mesoderm (Fig. S2D, E). These results indicated the pluripotency of piPSCs gradually vanished under F- and FD-treatments. The loss of pluripotency gene expression prompts us to check repression related epigenetic modification H3K9 methylation in these cells. The results revealed that the expression of *KDM3A/3B/3C/4C* genes was remarkably decreased and H3K9 methylation was gradually increased in F- and FD-treatments (Fig. 1C, D, F). Correspondingly, immunofluorescence showed that the level of H3K9me2/3 increased gradually, whereas the level of H3K4me3 which involved in the regulation of gene activation decreased (Fig. 1E, S2F). Further analysis demonstrated that the fluorescence intensity of H3K9me2/me3 at the marginal zone of clone was higher than that inside in the F-treatments (Fig. 1E), whereas the fluorescence intensity of H3K4me3 had an opposite trendency (Fig. S2G). This result was closely associated with the phenotype of cell marginal differentiation. These observations suggested that the increased H3K9 methylation in the F- and FD-treatments lead to cell differentiation. Moreover, the function of pluripotent factors may depend on the demethylation of H3K9me2/3 mediated by feeder.

### Feeder and pluripotent factors synergize to construct the pluripotency network

We attempted to gain insights into the molecular consequences that were triggered in response to the action of these treatments. We performed RNA sequencing to provide a snapshot of the transcriptional dynamics after the pluripotency changed. Transcriptional changes in both directions were observed in F- (272 up versus 443 down) and FD-treatments (1786 up versus 1144 down; Fig. 2A). Notably, down-regulated genes accounted for the majority in the differentially expressed genes (DEGs) in F-treatment (Fig. 2A). Given the significant increase of H3K9me2/3 in this treatment, feeders might promote the gene expression by facilitating the erasure of H3K9me2/3. However, more up-regulated transcription was observed in FD-treatment (Fig. 2A), suggesting that core pluripotent factors mainly inhibited transcriptional output to maintain pluripotency. Further analysis showed 200 of 272 up-regulated genes in the F-treatment were also up-regulated in FD-treatment, and 226 of 443 down-regulated genes in the F-treatment were also down-regulated in FD-treatment (Fig. 2B), indicating that these two treatments went through a consecutive process of differentiation. Additionally, the pluripotency-related genes and *KDM3A/3B/4C* showed a gradient descending trendency under F- and FD-treatments (Fig. 2C). This indicated feeder and exogenous pluripotency genes processed a complementary regulation during piPSCs maintenance.

**Fig. 2.**
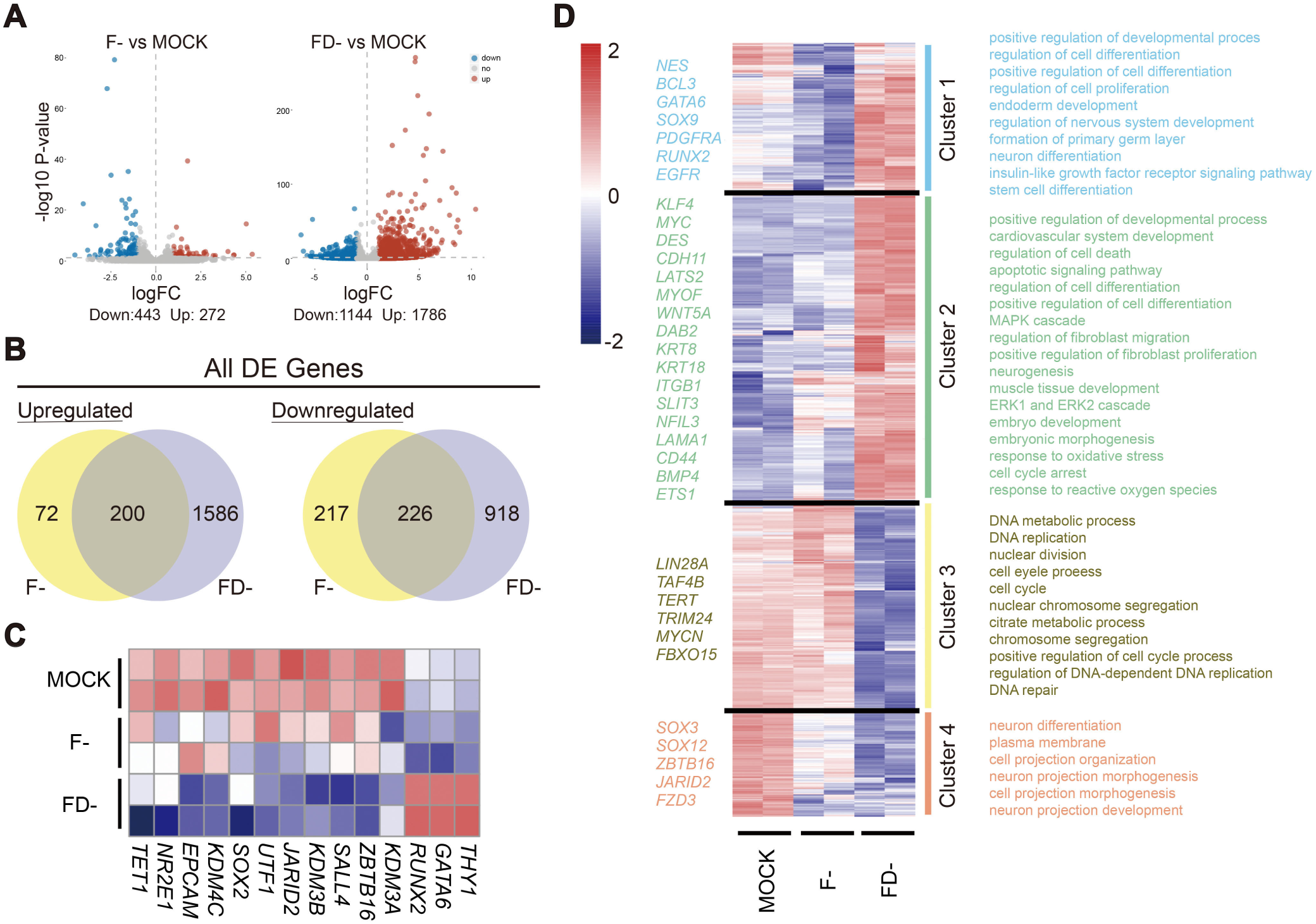
Feeder and exogenous genes cooperatively facilitate pluripotent transcriptional responses and inhibit the expression of differentiated related genes. **(A)** The volcano plot showing the number of up- or downregulated differentially expressed (DE) genes between F- and FD-conditions on day 5 determined by RNA-Seq. n = 2 independent experiments. **(B)** Venn diagrams showing the overlap of up- or downregulated differentially expressed (DE) transcripts between MOCK, F- and FD-conditions. **(C)** Heat map showing changes in the expression of pluripotency, fibroblast markers and epigenetic modification related genes in differentiating treatments obtained from RNA-Seq analysis. **(D)** Four clusters of differential expression highlight the key differences between MOCK, F- and FD-conditions. Gene ontology enrichment for each cluster is presented on the right. Represented genes of each cluster are listed on the left.

To further investigate this process, we applied cluster analysis for the DEGs. Gene ontology analysis of DEGs revealed significant enrichment in GO terms “positive regulation of developmental process”, “MAPK cascade”, “positive regulation of fibroblast proliferation”, and “cell cycle arrest” in response to F- and FD-treatments (Clusters 1 and 2; Fig. 2D). FD-responsive down-regulated genes were enriched for “neuronal differentiation”, “chromosome segregation”, and “DNA repair” (Clusters 3 and 4; Fig. 2D). These results indicated the DOX (exogenous genes) mainly inhibited the expression of differentiation-related genes and activated DNA repair-related genes to maintain the pluripotency of piPSCs. The cluster analysis also revealed that the feeder mainly promoted the expression of the SRY-related HMG-box (SOX) family of transcription factors (Clusters 1 and 4) to maintain the pluripotency of piPSCs which is consistent with feeder’s function reported before [25, 26]. Some transcription factors in SOX family (Cluster 4) were co-activated by feeder and exogenous genes, such as *SOX3* [27] (Fig. 2D). The result suggested that reprogramming factors cooperated with feeder to drive the pluripotency gene regulatory network.

### H3K9me2/3 prevents pluripotent transcription factors regulating on the target genes

To further investigate the role of H3K9 methylation on the maintenance of pluripotency, ChIP-seq of H3K9me2/3 was performed in MOCK and FD-groups. The ChIP-seq showed that the genome-wide level of H3K9me2/3 was significantly increased in FD-treatment (Fig. 3A, B). The binding sites of H3K9me2/3 were located mainly in the intergenic region and intron rather than promoter, and no significant difference was noted between the MOCK and FD-near the transcription start sites (Fig. S3A, B). These results suggested that H3K9me2/3 inhibited gene expression by acting on the other transcriptional regulatory regions other than promoters. Further analysis of de novo peaks revealed H3K9me2 and H3K9me3 only shared 2211 peaks in FD-treatment which account for 11% of the newly added peaks of H3K9me3. However, the analysis of the peak-associated genes showed that there were 226+474+2272 genes affected by H3K9me2 and H3K9me3 in FD-treatment in which only 226 genes were co-regulated, accounting for 19.4% of the H3K9me3 genes in this treatment (Fig. S3C). This result suggested that H3K9me3 cooperates with H3K9me2 to regulate the partial target gene under FD-treatment, while the binding sites are different. Combined analysis of ChIP-seq and RNA-seq of MOCK and FD-groups were performed to clarify the relationship between H3K9me2/3 and pluripotency maintenance. Considering that H3K9 methylation will silence gene expression, the associated genes were divided into four parts: the downregulated genes marked by FD-specified H3K9me2/3; the upregulated genes marked by MOCK specified H3K9me2/3. Further GO analysis of genes in Parts 1 and 3 reveal their enrichment in “positive regulation of fibroblast proliferation”, “positive regulation of apoptotic process”, “positive regulation of ERK1 and ERK2 cascade” and “positive regulation of epithelial to mesenchymal transition” (Fig. S3D). The genes in Parts 2 and 4 included several potential pluripotent genes, such as *NR6A1*(Fig. S3D) [28]. The result suggested that H3K9me2/3 participated not only in the inhibition of pluripotent genes during differentiation but also in the inhibition of somatic genes in the pluripotent state.

**Fig. 3.**
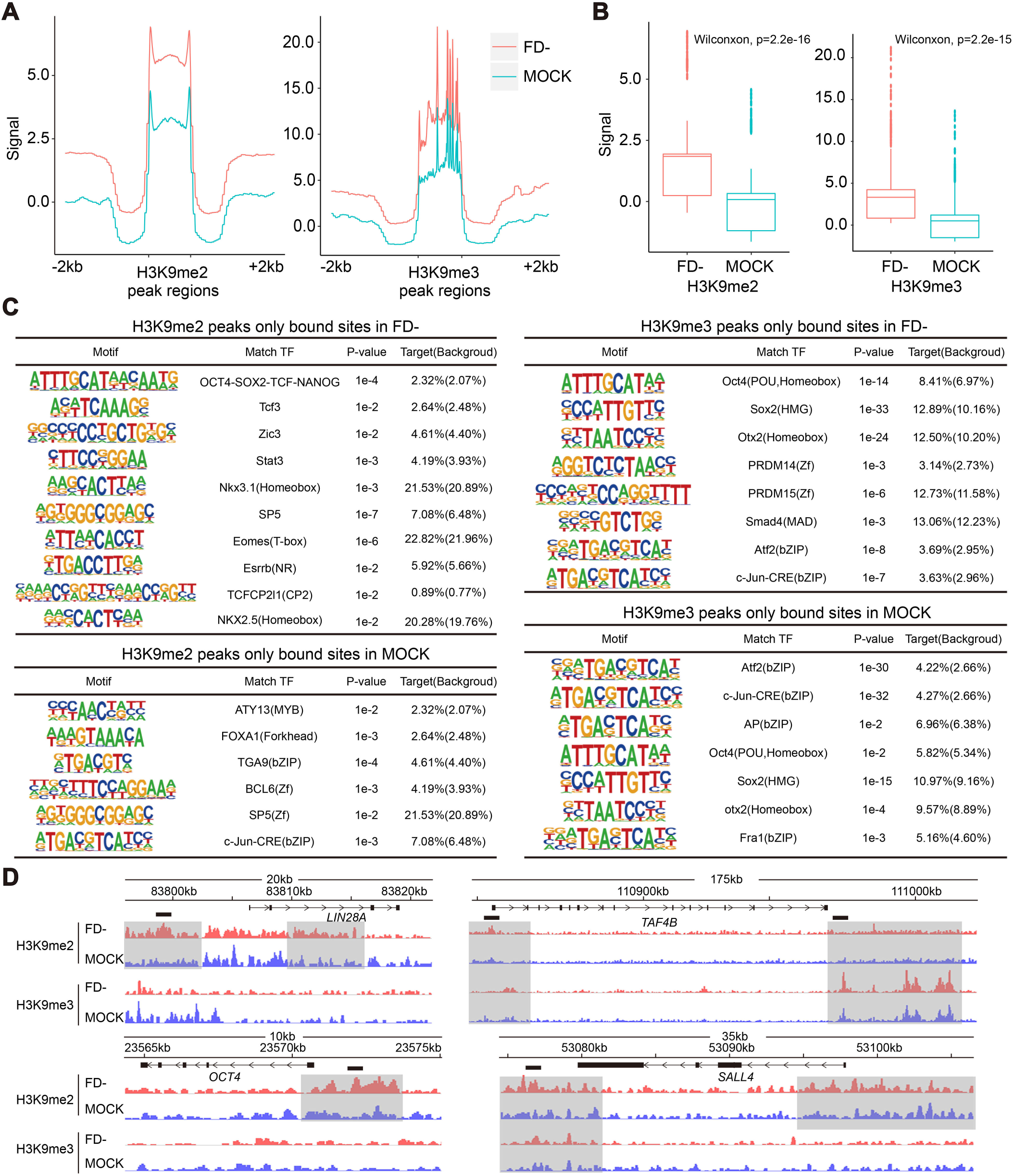
H3K9me2/3 prevents interaction between pluripotent transcription factors and target genes. **(A)** Metaplots of signal densities for H3K9me2/3 in MOCK and FD-treatments at all H3K9me2/3 bound sites. **(B)** Boxplots of global H3K9 methylation levels in MOCK and FD-treatments. **(C)** The motifs of H3K9me2/3 were identified at MOCK and FD-conditions. Last column: observed and expected motif frequencies (in parentheses). **(D)** Chromatin immunoprecipitation-sequencing (ChIP-seq) analysis of H3K9me2/3 marks at pluripotency gene loci in MOCK and FD-conditions using IGV. Gray bars represent the H3K9 methylation difference area in MOCK and FD-conditions. The size of the peak represents the degree of enrichment. The bold black line indicates the detection regions of ChIP-qPCR.

To further clarify the role of H3K9me2/3 in the maintenance of pluripotent state, motif analysis of H3K9me2/3 peaks was performed. Results revealed that H3K9me2 peaks only bound sites in FD-group contained pluripotent transcription factor binding sites, such as OCT4 and SOX2 (Fig. 3C). Although this finding is not clear on H3K9me3, more peaks have been found to contain pluripotency factors binding sites in FD-group compared to the MOCK group. This result implied that H3K9me2/3 prevented interaction between pluripotent transcription factors and their target genes, especially H3K9me2. To confirm this, we found a significant enrichment of H3K9me2 modification occurred in the upstream or downstream region of pluripotent genes *OCT4, LIN28A, TAF4B*, and *SALL4* in the FD-group (Fig. 3D), suggesting that these genes expression was suppressed (Fig. S3E, F). These results suggested that H3K9me2/3 represses gene expression by preventing the interaction between pluripotent transcription factors and target genes.

### Combined loss of KDM3A and KDM3B reveals critical roles of H3K9 demethylation in piPSCs maintenance

H3K9 methylation was unevenly distributed in chromosomes and regulated by KDM3A/B. However, the molecular mechanism remained unclear. To further directly investigate the role of H3K9 hypermethylation in the maintenance of piPSCs, we down-regulate the expressions of *KDM3A* and *KDM3B* by shRNA. Unexpectedly, the depletion of KDM3A only resulted in a slight decrease in the proliferation of piPSCs (Fig. S4A, B), and the global level of H3K9me2/3 was only slightly up-regulated (Fig. 4A, B). KDM3B loss not only significantly reduced the size of clones and proliferation rates but also decreased the degree of AP staining. However, the level of H3K9me1/2/3 remained unchanged (Fig. 4A, B). The results varied from previous studies on mESCs [6, 7]. KDM3A and KDM3B possess intrinsic H3K9 demethylating activity, so their functions may be complementary in the H3K9 demethylation. As expected, the co-depletion of KDM3A and KDM3B by shRNA induced a significant growth arrest and pluripotency loss in this group (Fig. 4A, B, S4A, B). Consistent to this, H3K9me2/3 was increased significantly (Fig. 4B). These results suggested that KDM3B could rescue the depletion of KDM3A, but KDM3A could not rescue that of KDM3B, suggesting that KDM3B might be involved in other biological processes.

**Fig. 4.**
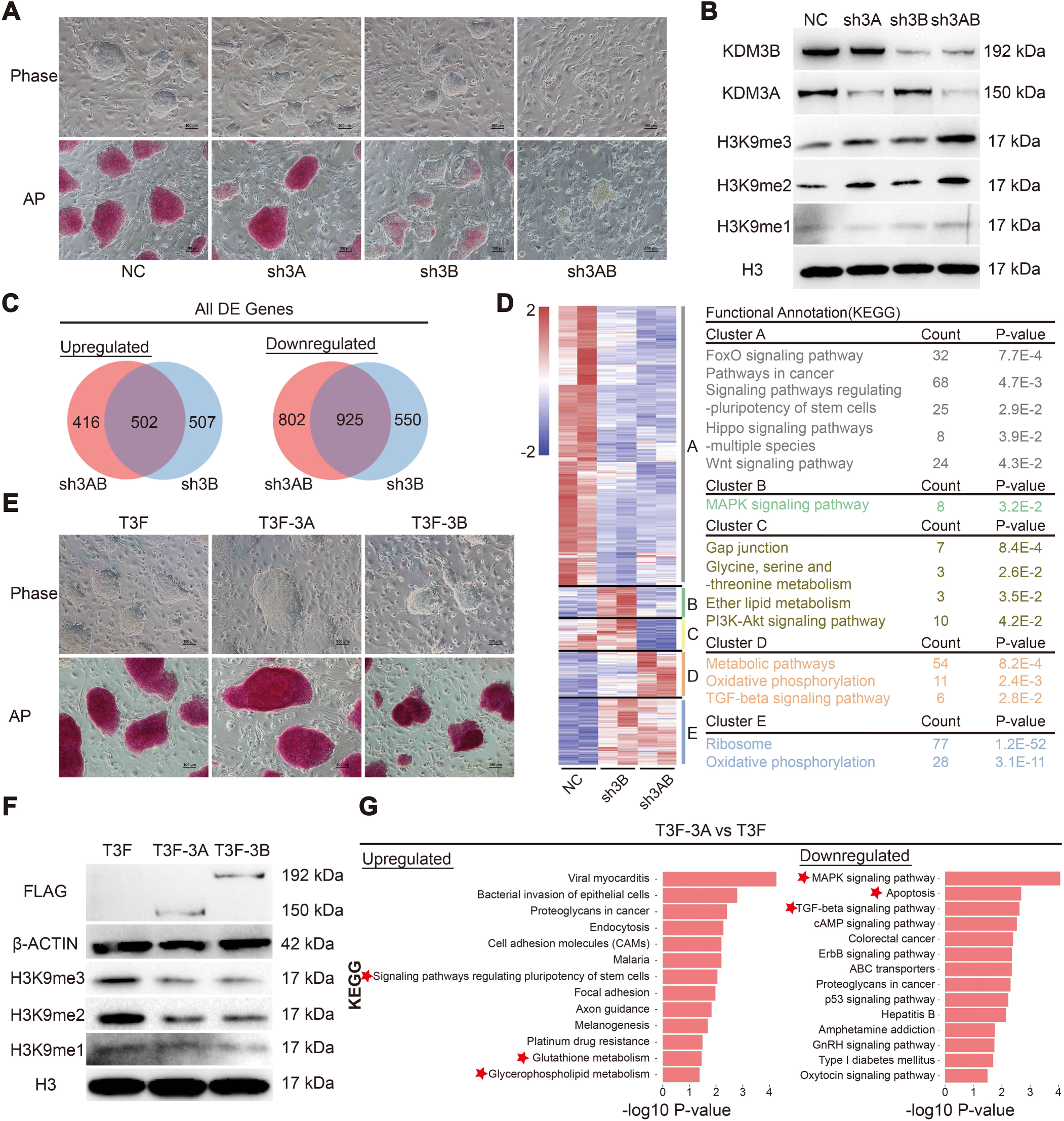
KDM3A and KDM3B synergize to maintain the pluripotency network in piPSCs. **(A)** Representative image of AP stained colonies after 5 days of clonal growth in the NC (Negative Control), sh3A(shKDM3A), sh3B(shKDM3B) and sh3AB (shKDM3A/B) cell lines. The experiments were performed three times. The scale bar represents 100 μm. **(B)** Representative Western-Blot of KDM3A, KDM3B and H3K9me1/2/3 after 5 days of culture in the normal condition. n = 3 independent experiments. **(C)** Venn diagrams showing the overlap of up- or downregulated differentially expressed (DE) transcripts between sh3B and sh3AB cell lines on day 5 determined by RNA-Seq. **(D)** five clusters of differential expression highlight the key differences between NC, sh3B and sh3AB cell lines. KEGG enrichment for each cluster is presented on the right. **(E)** Representative image of AP stained colonies after 5 days of clonal growth in the T3F (Empty vector), T3F-3A (overexpression of KDM3A) and T3F-3B (overexpression of KDM3A) cell lines. The experiments were performed three times. The scale bar represents 100 μm. **(F)** Representative Western-Blot of FLAG and H3K9me1/2/3 after 5 days of culture in the normal condition. n = 3 independent experiments. **(G)** KEGG enrichment of up- or downregulated differentially expressed (DE) genes in T3F-3A cell lines.

To molecularly characterize KDM3A/B function, the profiles of the transcriptomes of shRNA treated samples (shKDM3A, shKDM3B, and shKDM3A/B groups) were analyzed as well as empty vector control (NC) by RNA-sequencing. Although transcriptome did not change much in shKDM3A group (11 up versus 27 down), the gene expression in shKDM3B and shKDM3A/B was observed significantly changing with 1099 and 918 up-regulation; 1475 and 1727 down-regulation respectively (Fig. S4C). This finding indicates the depletion of KDM3A alone had no significant effect on the pluripotency of piPSCs. Further analysis revealed shKDM3B shared 50% (upregulated) and 63% (downregulated) dysregulated genes with shKDM3A/B group, explaining why KDM3B could rescue the lack of KDM3A (Fig. 4C). Of note, down-regulated genes (925 downregulated vs 502 upregulated) accounted for the majority in the DEGs of shKDM3B and shKDM3A/B treatments, which suggested KDM3A and KDM3B played transcriptional activation roles in the transcription regulation. The DEGs were then classified into five clusters (Fig. 4D). KEGG analysis was performed on each cluster to clarify the effects of KDM3A and KDM3B on the maintenance of piPSCs. Genes with functions in pathways of “signaling pathways regulating pluripotency of stem cells”, “Wnt signaling pathway”, and “Pathways in cancer” were significantly down-regulated in shKDM3B and shKDM3A/B groups (cluster A), whereas PI3K-Akt signaling pathway-associated genes were expressed at lower levels only in the shKDM3A/B group (cluster C). The regulation of associated genes on “oxidative phosphorylation” was up-regulated in the shKDM3B and shKDM3A/B groups (cluster E). Genes representing “TGF-beta signaling” and “metabolic pathways” were gradually up-regulated in the shKDM3B and shKDM3A/B groups, and these genes up-regulated more in shKDM3A/B group compared to shKDM3B only (cluster D). Unexpectedly, 268 genes representing MAPK signaling pathway were expressed at higher levels in the shKDM3B group, whereas the expression of these genes decreased after interfering with KDM3A expression on the basis of shKDM3B (cluster B; Fig. 4D). The transcriptome analysis results suggested that the depletion of KDM3A and KDM3B destroyed the pluripotency of piPSCs by decreasing expression of genes related to “Signaling pathways regulating pluripotency of stem cells”, “Wnt signaling pathway”, and “PI3K-Akt signaling pathway”. In addition, the piPSCs tended to differentiate and respond to oxidative stress, such as TGF-beta signaling pathway and oxidative phosphorylation, when depleted both KDM3A and KDM3B by shRNA (cluster D and E). These results suggested that erasure of H3K9me2/3 by KDM3A and KDM3B played a critical role in activating pluripotency related signaling pathways, such as Wnt signaling pathway, and PI3K-Akt signaling pathway.

### Histone demethylase KDM3A play a dominant role in maintaining pluripotency

Previous studies have demonstrated that the depletion of either KDM3A or KDM3B causes the increase of H3K9 methylation and loss of pluripotency in mESCs [6, 7]. However, a similar phenomenon only appeared in piPSCs when both KDM3A and KDM3B were down-regulated, implying a novel and complex regulatory mechanism in piPSCs. To further reveal the demethylation mechanism regulated by KDM3A/B, they were respectively overexpressed in piPSCs (Fig. 4E, F). The results showed that global H3K9me2/3 of piPSCs was remarkably decreased after overexpression of KDM3A or KDM3B (Fig. 4F, S4D). Differences in clone phenotypes were observed between T3F-KDM3A and T3F-KDM3B groups (Fig. 4E). Meanwhile, higher cell proliferation rate has been detected after KDM3A overexpression while KDM3B overexpression inhibits cell proliferation (Fig. S4E), suggesting that KDM3A and KDM3B were involved in different biological processes.

To understand the molecular mechanism of KDM3A and KDM3B in piPSCs regulation, we profiled the transcriptomes of T3F-KDM3A and T3F-KDM3B by RNA-seq. We found KDM3A (460 up versus 283 down) and KDM3B (378 up versus 258 down) had divergent transcriptional output (Fig. S4C). Only 128 up-regulated genes and 76 down-regulated overlapped genes shared by the two groups, accounting for 18% and 16.4% of the total number of differently expressed genes, respectively (Fig. S4F). The divergent transcriptional response again confirmed that KDM3A and KDM3B were involved in different biological processes. This finding explained why the depletion of KDM3B alone impairs the piPSCs pluripotency without altering global H3K9me2/3 modification. KEGG analysis was performed on the genes to clarify the effects of the overexpression of KDM3A and KDM3B on the maintenance of piPSCs. Genes with functions in pathways of “Signaling pathways regulating pluripotency of stem cells”, “glutathione metabolism”, and “glycerophospholipid metabolism” were significantly upregulated in T3F-3A group, whereas genes associated with MAPK, apoptosis, and TGF-beta signaling pathways were downregulated only in T3F-3A group (Fig. 4G). Nonetheless, overexpression of KDM3B enriched pathways unrelated to pluripotency (Fig. S4G). We also found naive pluripotent gene *TBX3* [29, 30] and *WNT* ligands upregulated while the expression of *JUN* which inhibiting reprogramming downregulated after overexpression of KDM3A [2] (Fig. S4H). However, the overexpression of KDM3B does not have the same effect. Together, our results suggested that the KDM3A maintained the pluripotent network of piPSCs by activating the expression of genes of signaling pathways regulating pluripotency as well as inhibiting genes associated with MAPK, apoptosis, and TGF-beta signaling pathways. The co-depletion of KDM3A and KDM3B influenced the pluripotency of piPSCs, but only overexpression of KDM3A could promote the pluripotency, suggesting KDM3A play a dominant role in maintaining pluripotency by H3K9 demethylation.

### OCT4 and SOX2 are the core factors for the maintenance of piPSCs

Considering that the downregulation of OSKM also leads to the increase of H3K9 methylation under F-condition, it is necessary to further investigate the role of core pluripotent genes in the maintenance of pluripotency. After removing DOX, the proliferation of piPSCs was significantly decreased, and the classical ESC like clonal morphology gradually destroyed (Fig. S1F, 5A). The expressions of endogenous *OCT4* and *SOX2* were significantly down-regulated, whereas those of endogenous *KLF4* and *c-MYC* were significantly up-regulated in FD-treatment (Fig. 1B). Therefore, four pluripotency genes OSKM may have different roles in piPSCs maintenance. Next, we try to determine single pluripotency gene’s function by using a lentivirus overexpression system carried EF1 promoter (Fig. S5A, B, C). The results showed that piPSCs could maintain clonal morphology under DOX-condition when OCT4 or c-MYC was complemented (Fig. S5A). However, the AP staining of c-MYC group gradually presented negatively as the passage times increased under the DOX-condition (Fig. S5D). Given the role of SOX2 may depend on OCT4, we then try to overexpress both factors in piPSCs to explore whether O/S can recapitulate exogenous OSKM function to maintain pluripotency. After overexpressed O/S (EF1α-OCT4-P2A-SOX2) or OCT4 (EF1α-3×FLAG-OCT4) alone, cell lines can be expanded more than 20 passages, and still maintain the phenotype without differentiation (Fig. 5A). The proliferation rate is similar when compared OS co-overexpression cell line with the control, however, the OCT4 overexpressing line is lower than the control (Fig. S5E). Further analysis showed that the expression of total *SOX2* was positively correlated with *LIN28A* expression level and negatively correlated with *NANOG* and *c-MYC* (Fig. 5B). These results indicated that SOX2 was also a core transcription factor that regulated the proliferation of piPSCs, and its function depended on OCT4.

**Fig. 5.**
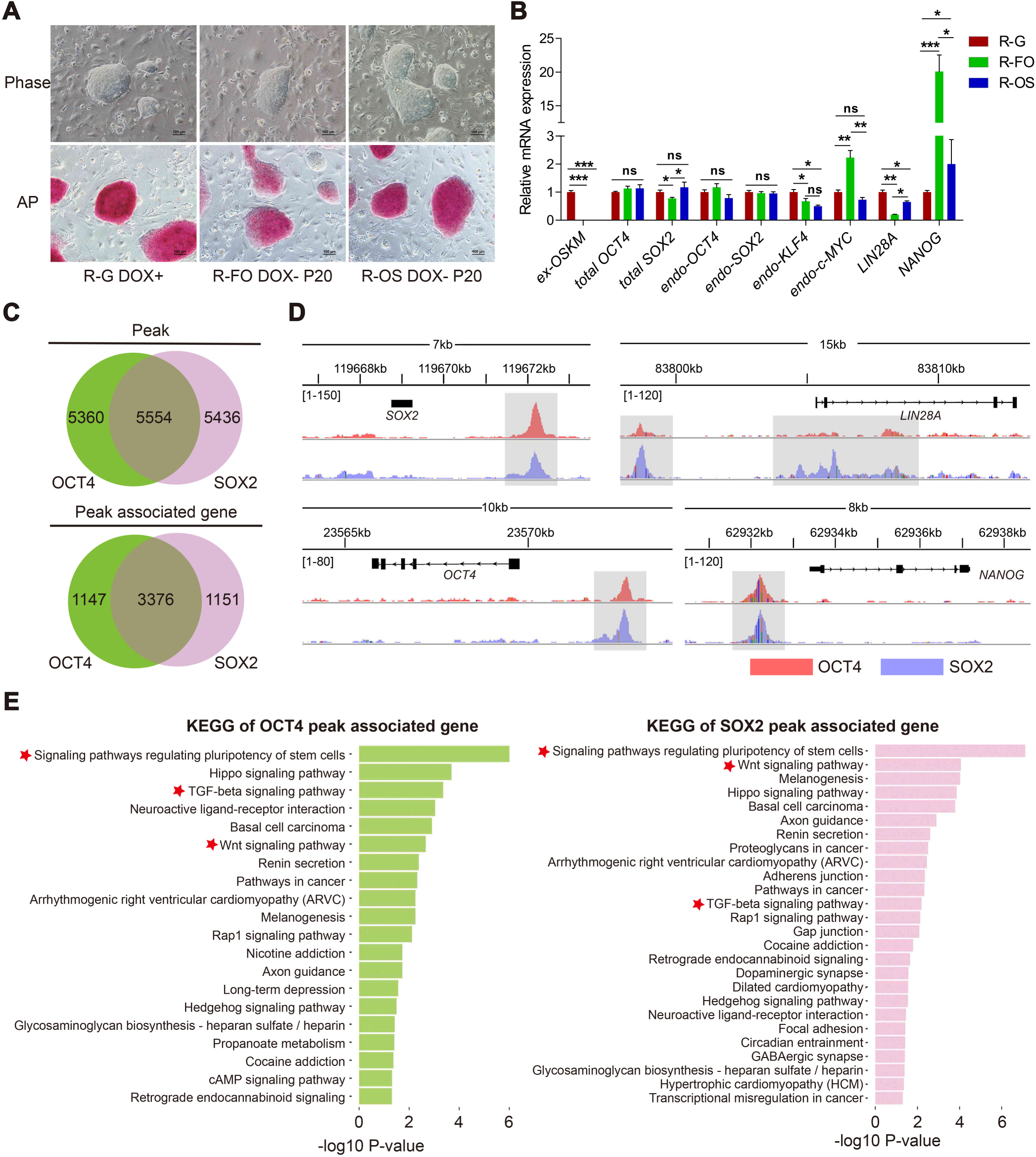
OCT4 and SOX2 was the core factors for the Maintenance of piPSCs. **(A)** Representative image of AP stained colonies after 5 days of clonal growth in the R-G (Rosa26-EF1α-GFP, Negative Control), R-FO (Rosa26-EF1α-3×flag-OCT4) and R-OS(Rosa26-EF1α-OCT4-P2A-SOX2) cell lines. The DOX+/- represents the presence/absence of Dox in medium. The scale bar represents 100 μm. **(B)** RT–qPCR analysis of the exogenous reprogramming factors and endogenous pluripotent genes in the R-G, R-FO and R-OS treatments. The relative expression levels were normalized to β-actin. Data represent the mean ± s.d.; n = 3 independent experiments. **(C)** Venn diagrams showing the overlap of OCT4 and SOX2 peak and peak associated gene determined by ChIP-Seq. **(D)** Chromatin immunoprecipitation-sequencing (ChIP-seq) analysis of OCT4 and SOX2 at pluripotency gene loci in normal condition using IGV. Gray bars represent the OCT4 and SOX2 binding area. The size of the peak represents the degree of enrichment. **(E)** KEGG enrichment of OCT4 and SOX2 peak associated gene.

ChIP-seq of OCT4 and SOX2 were performed to analyze their mechanism of transcriptional regulation. The results showed that the binding sites of OCT4 and SOX2 were mainly located in the intergenic region and intron rather than promoter, suggesting that they activated gene expression by acting on the distal enhancer region (Fig. S5F). It is noteworthy that the OCT4 shares about 50% (5554) peaks with SOX2, which accounts for 74.6% (3376/4523) of the genes OCT4 target (Fig. 5C), supporting interaction between OCT4 and SOX2 during binding to maintain gene expression. These findings were similar to the results reported in mouse [1, 31, 32]. Motif analysis indicating OCT4 and SOX2 were of high similarity (Fig. S5G), which is consistent with the peak binding analysis between the two proteins. Therefore, we performed a bimolecular fluorescence complementary test to directly investigate the proteins interacting with OCT4 and SOX2 (Fig. S7B).

To reveal why OCT4 and SOX2 were necessary for piPSCs, we further analyzed their peaks in pluripotency genes *OCT4, SOX2, LIN28A*, and *NANOG*. ChIP-seq showed that all the genes were co-occupied by OCT4 and SOX2 (Fig. 5D). We further validated the regulation by double luciferase test and found all the four genes can be activated by O/S, and *LIN28A* showing significantly higher luciferase activity (13-fold change) than other genes (Fig. S5H). KEGG analysis also revealed OCT4 and SOX2 targets are mainly enriched in the pathway related to “Wnt/β-catenin signaling pathway”, “TGF-β signaling pathway”, and “signaling pathway regulating pluripotency of stem cells” (Fig. 5E). These results indicated OCT4 and SOX2 formed heteromeric dimers to maintain the porcine pluripotent regulatory network.

### O/S and KDM3A/3B synergize to construct the porcine pluripotent networks

Previous studies in mES have demonstrated the direct activation of KDM3A/3B by OCT4 [6, 8]. To determine if this regulation link still exists in piPSCs, we overexpressed OCT4 and SOX2 in piPSCs (Fig. S6A). The results showed that the expression of *KDM3A* and *KDM3B* were not affected, and the promoter activity of *KDM3A* and *KDM3B* cannot be activated by OCT4 and SOX2 (Fig. S6B, C). In addition, the expression of endogenous *OCT4* and *SOX2* did not further increase significantly when KDM3A or KDM3B were overexpressed (Fig. S6D). Given the dependence between transcription factors O/S and hypomethylation of H3K9 in this study (Fig. 3C), the relationship between them might be a cooperator. To further explore the physiological relevance of our findings that OCT4 and SOX2 maintain piPSCs pluripotency is dependent on H3K9 hypomethylation, we ask how many OCT4 and SOX2 peaks co-occupied by H3K9me2/3. Finally, we found about 20% OCT4 (18%, 1984/10817) and SOX2 (22%, 2346/10870) peaks modified by H3K9me2 while fewer peaks have been found modified by H3K9me3 in FD-condition (Fig. 6A). Furthermore, we analyzed the change of H3K9me2/3 in OCT4 and SOX2 binding sites. Previous results showed that there was no significant difference near the transcription start sites (Fig. S3B), but the level of H3K9me2 at the OCT4 and SOX2 binding sites increased significantly in FD-group, indicating that H3K9 methylation directly prevented the function of OCT4/SOX2 (Fig. 6B, C). The KEGG enrichment of peak-associated genes shared by OCT4/SOX2 and H3K9me2 showed that the genes were enriched in pathways like Wnt/β-catenin signaling pathway which is important for pluripotency of stem cells regulation (Fig. 6D). This result indicated that OCT4 and SOX2 drove critical pluripotent pathway depending on the erasure of H3K9me2. In addition, ATAC-seq of MOCK and FD-groups also showed that with the increase of H3K9me2 in the O/S binding sites, the accessibility of these sites in the FD-group decreased significantly (Fig. 6E). Motif analysis further showed that the proportion of pluripotency related motifs (OCT4/SOX2) was decreased, while the proportion of somatic transcription factors (Fra1/Jun-AP1/AP1) were increased in FD-treatment, indicating the OCT4/SOX2 binding patterns change in this treatment (Fig. S6E). Interestingly, conjoint analysis between the RNA-seq of T3F-3A and ChIP-seq of O/S not only reconfirmed the above results but also found KDM3A was involved in the inhibition of O/S-mediated TGF-β and MAPK signaling pathways (Fig. S6F, G). ChIP-seq analysis further validates OCT4 and SOX2 binding to the enhancer and promoter regions of pluripotent genes *OCT4, LIN28A, NANOG*, and Wnt receptor proteins *FZD8* in piPSCs, but these binding regions showed increased H3K9me2 in the FD-treatment, that is, the binding sites of OCT4 and SOX2 were closed (Fig. 6F). These results suggested that OCT4 and SOX2 activated key pluripotent genes depending on the erasure of H3K9me2.

**Fig. 6.**
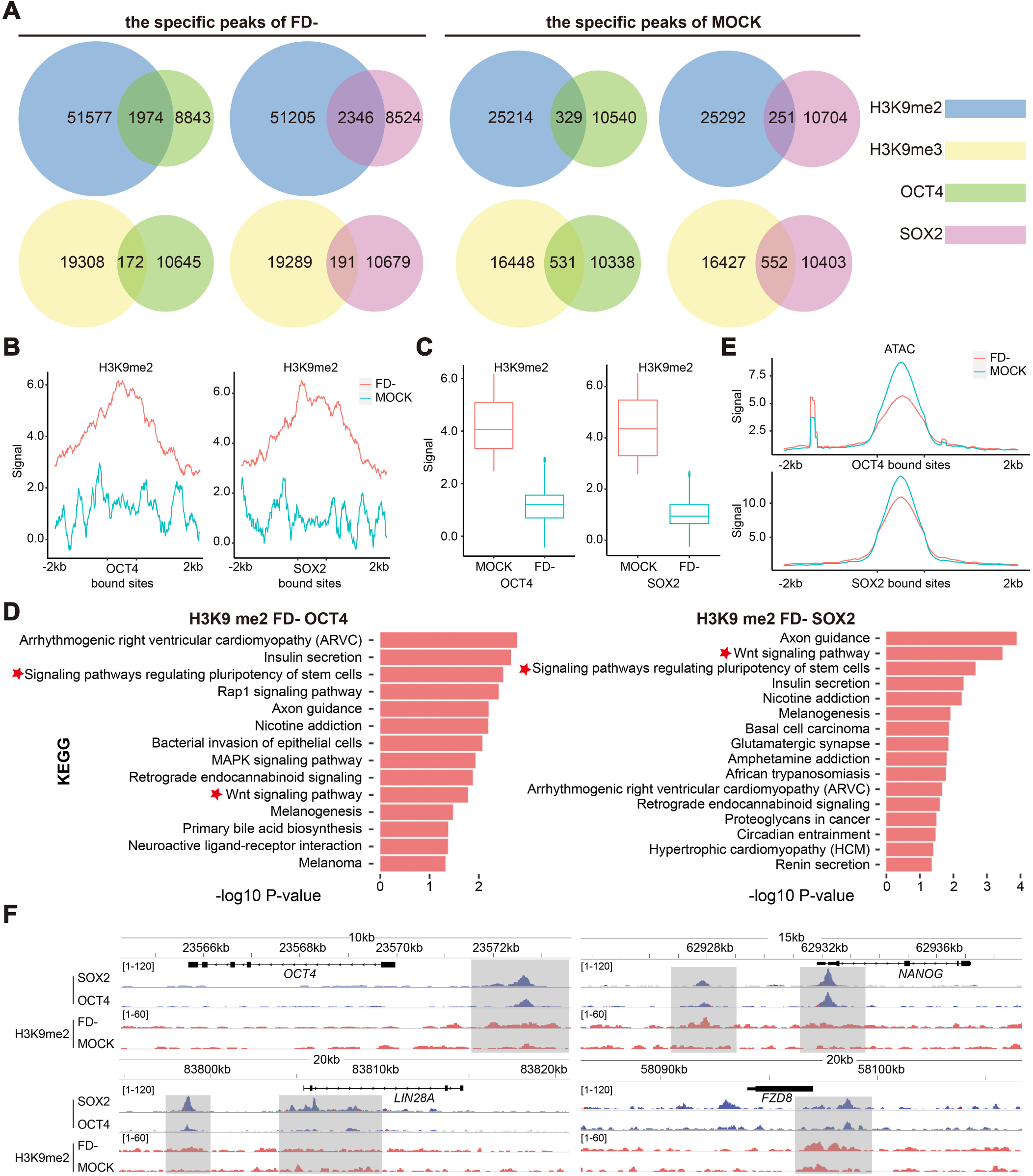
The demethylation of H3K9me2 is essential for O/S to activate pluripotent networks. **(A)** Intersection of OCT4 or SOX2 binding sites and the specific H3K9me2/3 peaks of MOCK and FD-. The number of peaks for each part is given in the Venn diagram. Blue for H3K9me2 peaks, yellow for H3K9me3 peaks, green for oct4 peaks and red for SOX2 peaks. **(B)** Metaplots of signal densities for H3K9me2 in MOCK and FD-treatments at all OCT4 and SOX2 bound sites. **(C)** Boxplots of H3K9me2 levels in MOCK and FD-treatments at all OCT4 and SOX2 bound sites. **(D)** KEGG enrichment of the intersection of OCT4 or SOX2 binding sites and the specific H3K9me2 peaks in FD-associated gene. The red marks represent major pluripotent regulatory pathways. **(E)** Metaplots of signal densities for ATAC-seq in MOCK and FD-treatments at all OCT4 and SOX2 bound sites. **(F)** The chromatin immunoprecipitation-sequencing (ChIP-seq) analysis of OCT4 and SOX2 at pluripotency gene loci and H3K9me2/3 marks at pluripotency gene loci in MOCK and FD-conditions using IGV. The size of the peak represents the degree of enrichment.

To further dissect the demethylation mechanism underlying KDM3A and KDM3B in the O/S binding sites, we applied IP-MS to directly investigate the proteins interacted to KDM3A and KDM3B under normal condition (Fig. 7A, B, S7A). The results showed an interaction between KDM3A and KDM3B. Additional studies were conducted to verify the direct binding between KDM3A and KDM3B using a bimolecular fluorescence complementary (BiFC) technique (Fig. S7B). GO analysis was performed on the proteins belonging to each part to clarify the different effects of KDM3A and KDM3B on the maintenance of piPSCs. Partners shared by KDM3A and KDM3B were enriched with functions in “DNA ligation involved in DNA repair”, “histone H3-K9 demethylation”, and “DNA topological change”. KDM3A specific interacting proteins were enriched with functions in “positive regulation of telomerase activity”. The specific KDM3B-interacting proteins were enriched with functions in “positive regulation of Wnt protein secretion” and “glycolytic process” (Fig. 7C). These results suggested that KDM3A and KDM3B co-regulated gene activation processes in the chromatin level but also individually involved in different biological processes.

**Fig. 7.**
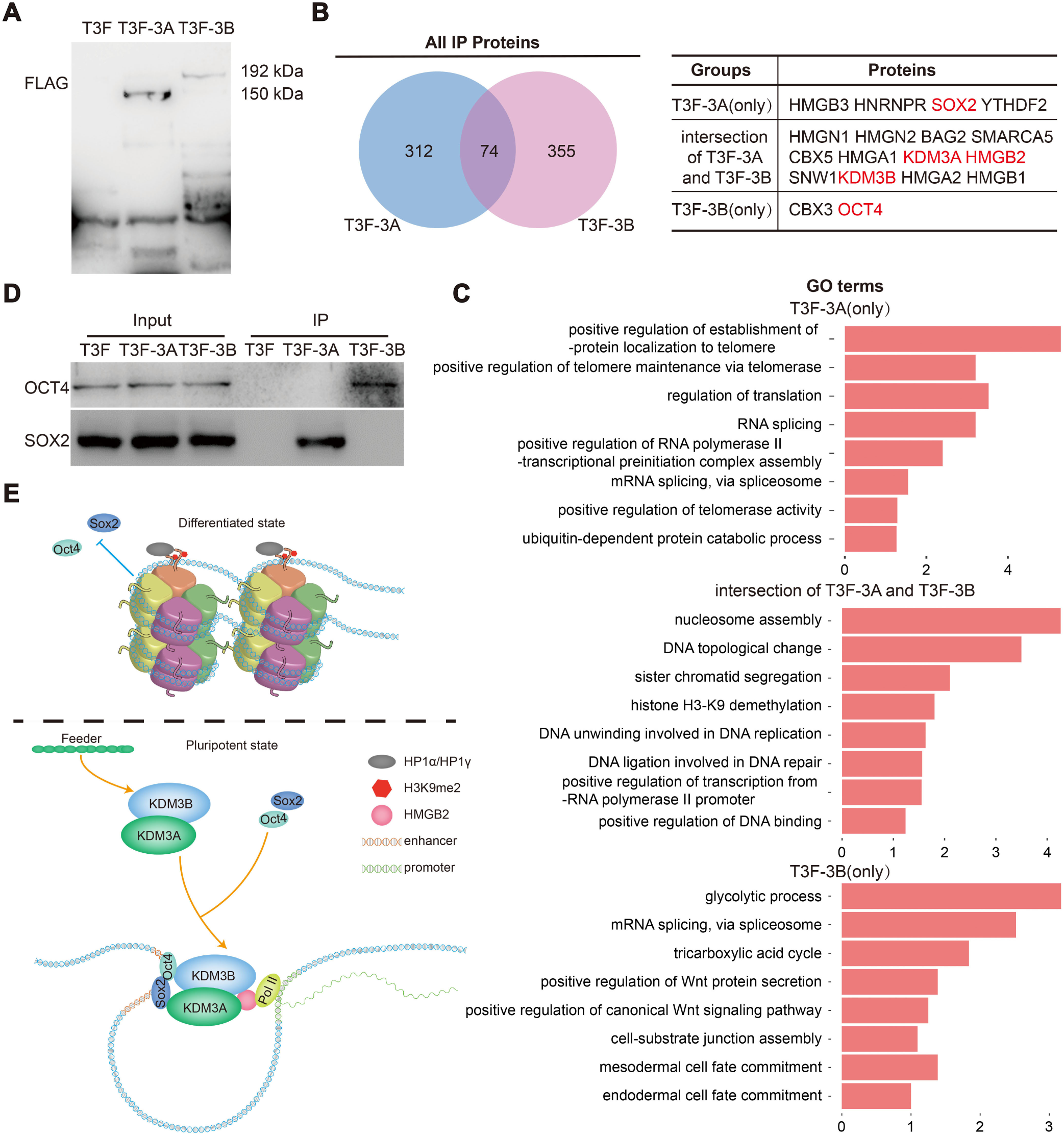
Cooperative binding of core transcription factors and KDM3A/B to maintain pluripotency. **(A)** Immunoprecipitation of KDM3A and KDM3B were validated using western blot. piPSCs expressing 3×FLAG-KDM3A or 3×FLAG-KDM3B were used to purify KDM3A or KDM3B-associated proteins. The KDM3A or KDM3B-associated proteins were also detected and identified by mass spectrometry. **(B)** Venn diagrams showing the overlap of KDM3A and KDM3B associated proteins determined by mass spectrometry. The table lists some representative proteins of each section. **(C)** Gene ontology enrichment of KDM3A or KDM3B-associated proteins. **(D)** The correlation between KDM3A, KDM3B, SOX2 and OCT4 were validated by western blotting. **(E)** Model for action of core transcription factors and KDM3A/B during pluripotency maintenance.

Further analysis of the IP-MS revealed that KDM3A interacted with SOX2, whereas KDM3B interacted with OCT4, which were further confirmed using IP-WB (Immunoprecipitation combined with western blot) (Fig. 7B, D). A large number of high mobility group (HMG) proteins involved in chromatin remodeling [33, 34] were significantly enriched in the intersection of the IP-MS of KDM3A and KDM3B (Fig. 7B). BiFC test again demonstrated the direct interaction between OCT4, SOX2, KDM3A, and KDM3B (Fig. S7B). These results suggested that OCT4/SOX2, KDM3A/KDM3B, and HMG proteins formed a transcription complex to activate downstream genes (Fig. 7E). This new mechanism of pluripotency maintenance is important for the establishment of porcine naïve-state PSCs.

## Discussion

The deconstruction of pluripotency gene regulatory network is a challenging and urgent issue need solving to obtain bona fide porcine pluripotent cells across the globe. In this study, we used the piPSCs generated by TetO-inducible system to gain a clear description of how the porcine epigenetic pluripotency network was maintained. Our results demonstrated that OCT4 and SOX2 cooperate with KDM3A/KDM3B to support the complex formation of super-transcription drivers to porcine pluripotency network (Fig. 7E).

KDM3A/KDM3B-meditated H3K9 demethylation plays a critical role in early mouse embryogenesis and mES maintenance [6-8]. However, no detailed information can be found on H3K9 methylation regulating porcine embryogenesis and ESC maintenance. In addition, the rules of KDM3A and KDM3B in H3K9 demethylation remained unclear. Our results demonstrate that KDM3A and KDM3B form a synergistic complex to perform demethylation, and only suppressing both of them will achieve global H3K9 hypermethylation. The conventional view is that KDM3A and KDM3B may function in a redundant manner [8-10]. However, our study also suggests that KDM3A and KDM3B are novel cooperative mechanisms in the erasure of H3K9 methylation. The single-cell sequencing of porcine embryo shows that KDM3A is highly expressed in Morula, ICM, and Epiblast, but KDM3B is only specifically expressed in Epiblast (Fig. S7C). The expression pattern was similar to *SOX2*, suggesting that a similar interaction mechanism might exist in Epiblast [35]. Our results also reveal that the overexpression of KDM3A can significantly improve the proliferation and anti-apoptosis ability of pluripotent stem cells, but that of KDM3B leads to a decline in cell proliferation. The results suggest that KDM3A is the preferred choice for the H3K9 demethylation and excessive KDM3B affects pluripotency by promoting other biological processes under normal condition. Our study provides a novel mechanism in which histone demethylase performs H3K9 demethylation in porcine pluripotent stem cells.

Porcine pluripotent stem cells, including embryonic stem cells (ESCs) and iPSCs, are facing a very serious problem [36]. At present, piPSCs can be obtained by ectopic expression of defined pluripotent transcription factors [24, 37, 38]. However, the maintenance of pluripotent state depends on the continuous expression of exogenous genes, and endogenous pluripotent genes are not fully activated [22-24, 37-41]. Therefore, most of piPSCs also fail to contribute to chimeras with exceptions that are only detected by genomic PCR analysis [22, 23]. Porcine embryonic stem cell-like cells (pESLCs) exhibited some features of pluripotency, but these cells could not maintain self-renewal for a long time [42-44] [45]. The generation of pESCs and acquisition of imperfect piPSCs always fail because the porcine endogenous pluripotent network cannot be maintained stably.

OCT4, which is a core factor during reprogramming, is often used as a marker for reprogramming cells to reach iPSC state [37, 46, 47]. Numerous studies have shown that the depletion of OCT4 causes pluripotent stem cells to gradually lose their pluripotency [32, 48]. Our study demonstrates that the deficiency of endogenous OCT4 expression is the main reason porcine pluripotent stem cells cannot be maintained for a long time. Several studies on pig early embryos have indicated that OCT4 is generic expressed in ICM and TE, which differs from that in humans and mice (Fig. S7C) [35]. The regulatory mechanism of endogenous OCT4 in pigs may be unique. Although our study did not further increase endogenous OCT4 by changing the culture system, we demonstrated that low expression of OCT4 is associated with high levels of H3K9me2 in its promoter and enhancer region (Fig. 3D). We speculate that H3K9me2 level must be decreased in OCT4 promoter region during the establishment of porcine ESCs.

SOX2, a molecular marker specifically expressed in pig ICM [49], cooperates with OCT4 to activate downstream pluripotent genes [31, 32]. According to our results, OCT4 and SOX2 together directly activate many pluripotency related pathways and regulate each other, and this finding is similar to that in other reports [1, 31, 32]. However, differences are noted in downstream pathways and genes, especially the TGF-β signaling pathway, which must be directly inhibited by O/S in piPSCs. The IP-MS of KDM3A and KDM3B shows that both bind a large number of proteins associated with chromatin remodeling, such as HMGB2. Our further study demonstrates that SOX2 and OCT4 directly bind to KDM3A and KDM3B, respectively, suggesting that the SOX2/OCT4 recruit KDM3A/KDM3B and chromatin remodeling proteins form a transcriptional complex to drive the pluripotency gene regulatory network. This novel mechanism is similar to the previously reported super enhancer. Understanding the mechanistic basis for forming the complex between pluripotent and non-pluripotent cells reveals further insights into the nature of the pluripotent state.

## Material and Methods

### Statement of Ethics

All animal experiments were performed in strict accordance with the Guide for the Care and Use of Laboratory Animals (Ministry of Science and Technology of the People’s Republic of China, Policy No. 2006398) and were approved by the Animal Care and Use Center of the Northwest A & F University.

### Cell culture

Porcine iPSCs was cultured on feeder (MEF, mouse embryonic fibroblasts) and maintained in LB2i medium, consisting of DMEM (Hyclone, USA) supplemented with 15% FBS (VISTECH, New Zealand), 0.1 mM NEAA (Gibco, USA), 1 mM L-glutaMAX (Gibco, USA), 10 ng/ml LIF (Sino Biological, 14890 -HNAE), 10 ng/ml bFGF (Sino Biological,10014-HNAE), 0.1 mM β-mercaptoethanol (Sigma Aldrich, M3148, USA), 3 µM CHIR99021 (MCE, HY-10182), 2 µM SB431542 (Selleck, S1067), 4 µg/ml doxycycline (Sigma Aldrich, D9891), 100 units/ml penicillin and 100 μg/ml streptomycin. piPSCs were passaged using TrypLE™ Select (Invitrogen, USA) into a single cell at 2×10^4^ cells per 12-well plate every 5–6 days. Differentiation was performed in differentiation medium: DMEM (Hyclone, USA) supplemented with 15% FBS, 0.1mM NEAA, 1 mM L-glutaMAX, 0.1 mM β-mercaptoethanol, 100 units/ml penicillin and 100 μg/ml streptomycin. HEK293T and MEFs were cultured in DMEM supplemented with 10% FBS and 100 units/ml penicillin, 100 μg/ml streptomycin [21].

### Plasmids and cloning

All lentivirus backbones vectors were derived from pCDH-CMV-MCS-EF1-GreenPuro (CD513B-1, SBI, Mountain View, CA, USA) by Seamless Cloning and Assembly Kit (Novoprotein, China). The following vectors used in this study were constructed using the same strategy and its information is shown in Table S1.

### shRNA vectors

shRNAs were designed using the online shRNA design tool from Invitrogen (http://rnaidesigner.thermofisher.com/rnaiexpress/setOption.do?designOption=shrna&pid=5230999276329139407/) with default parameters. And the potential off-target effects were filtered out using BLAST against the porcine genome.

Sense and antisense oligonucleotides of the shRNA duplex were synthesized and cloned into an in-house designed pCDH-U6-MCS-EF1-GFP-T2A-PURO or pCDH-U6-MCS-EF1-GFP-T2A-mCherry Lentivector. All shRNAs were tested for each gene in piPSCs and knockdown efficiencies of 70% or more were used in this study. The vector and sequence information of shRNAs used in this study is shown in Table S1 and S2.

### Overexpression vectors

The p-O/S/K/M/L(*OCT4/SOX2/KLF4/c-MYC/LIN28A*) gene were PCR-amplified from porcine blastocysts and subcloned into an in-house pCDH-EF1-MCS-T2A-PURO vectors. The 3×FLAG-OCT4, 3×FLAG-SOX2, 3×FLAG-KDM3A and 3×FLAG-KDM3B were generated by PCR and subcloned in pCDH-TetO-3XFLAG-MCS-T2A-PURO. The Rosa26-EF1α-3×FLAG-OCT4, and Rosa26-EF1α-OCT4-P2A-SOX2 were generated by PCR and subcloned in Rosa26-EF1α-MCS-T2A-PURO. Coding regions of all overexpression vectors were verified by sequencing. Subsequently, these overexpression vectors were transfected into HEK293T cell and identified by Western blot. The vector information of these genes used in this study is shown in Table S1.

### BiFC vectors

The *OCT4/SOX2/KDM3A/KDM3B/WNT2* gene were generated by PCR and subcloned in pBiFC-VC155 and pBiFC-VN173. Coding regions of all overexpression vectors were verified by sequencing. The vector information of these genes used in this study is shown in Table S1.

### Luciferase vectors

To verify the ChIP-seq of OCT4 and SOX2, the porcine target gene promoter and enhancer were amplified by PCR from total genomic DNA extracted from piPSCs and subcloned into an in-house PGL3-basic vector (Promega, E1751) by Seamless Cloning. The detailed information of these Luciferase vectors is shown in Table S1.

### Lentivirus packaging and transduction

HEK293T cells were seeded onto a 6-well plate and grown to 70–80% confluence. Then the lentivirus backbone and package vector (pVSV-G and psPAX2) were transfected into HEK293T cells using PEI (polyethyleneimine, sigma). For transfection of per well, 1 µg pVSV-G, 1µg psPAX2 and 2 µg lentivirus backbone vector were diluted in 200 µl optiMEM and vortexed. 12 µl PEI (1mg/ml) was added to the plasmid mix, vortexed, incubated for 15 min at room temperature then added to cells. After 12 h, the medium was replaced with fresh medium added lipid and cells were maintained for 48 −72 h. Lentivirus (culture supernate) was collected, filtered through 0.45 µm filter to remove debris. For the lentiviral transduction, 2×10^4^ cells were plated on MEF-coated 12 well plate per well and allowed to attach overnight. Virus and fresh media were added at a ratio of 1:1 supplemented with 4 µg/ml Polybrene into this well. The Cells were incubated with mixed media overnight, washed with PBS and replaced with fresh medium. After 1 week of culture, stably infected colonies were selected with puromycin (10 µg/ml) for 24 h, and viral titer was calculated by counting the GFP-positive colonies.

### AP staining

Cells were fixed with 4% paraformaldehyde in PBS (pH 7.4) for 15 min at room temperature, washed twice using ice-cold PBS and developed with AST Fast Red TR and α-Naphthol AS-MX Phosphate (Sigma Aldrich) according to the manufacturer’s instructions. Then the cells were incubated with the mixture (1.0 mg/ml Fast Red TR, 0.4 mg/ml Naphthol AS-MX in 0.1 M Tris-HCL Buffer) at room temperature. After 20 min, the AP-positive iPS colonies showed in red color. The images were collected by a Nikon phase contrast microscope.

### RNA extraction, Reverse transcription, and Quantitative real-time PCR

Total RNA was extracted by RNAiso Plus (Takara, 9109) and purified with the guanidine isothiocyanate - phenol chloroform. Reverse transcription was performed using Fast Quant RT Kit (TIANGEN, KR106). Fluorescence quantitative PCR analyses of all samples were performed using a Bio-Rad CFX96 and SYBR green master mix (TIANGEN, FP215). Semi-quantitative PCR reactions were performed for 30 cycles by 2×TSINGKE Master Mix (TSINGKE, TSE004). The primer information used in this study is provided in Table S3. The expression of target gene was normalized against transfection of control vector. Data of Q-PCR are calculated by Bio-Rad software CFX3.1 and derived from three independent experiments.

### Western blot

The cells were digested by TrypLE™ Select and then transferred immediately to a 1.5 ml tube on ice. The cell suspension was then centrifuged at 5000g for 3 min and supernatant was discarded. The cell sediment was lysed by RIPA buffer (Beyotime, P0013B) for 30 min on ice, added to 5× SDS-PAGE loading buffer (GENSHARE G, JC-PE007), and heated at 100 °C for 5 min, then loaded onto 8-12% SDS-PAGE gel. The SDS-PAGE gels were run at 100V for 1.5 h and transferred to a PVDF membrane by semidry electrophoretic transfer (Bio-Rad) for 45 min at 15 V. The transferred membrane was blocked with 8% skim milk) at room temperature for 2 h, and then incubated with the primary antibody in TBS-T buffer (20 mM Tris/HCl pH 8.0, 150 mM NaCl, 0.05% Tween 20) at 4 °C overnight. After washing three times with TBS-T buffer, the blot was incubated with secondary antibody at 37°C for 1h, then washed three times. Enhanced chemiluminescent substrate (Biodragon, BF06053-500) was used to detect the HRP signal and the western blot images were collected using the Chemiluminescent Imaging System (ZY058176, Tanon-4200, China). The information of antibodies used in this study was listed in Table S4.

### Immunofluorescence microscopy

The cells were fixed with 4% paraformaldehyde in PBS (pH 7.4) for 15 min at room temperature. Fixed cells were washed twice using ice-cold PBS, permeabilized with 0.1% Triton X-100 in PBS for 10 min, and subsequently blocked for 2 h at room temperature in PBS containing 5% FBS. The cells were incubated with blocking buffer containing primary antibodies at 4°C overnight. The secondary antibodies were diluted in a blocking buffer and incubated at 37°C for 1h. After washing with PBS for three times, the nuclei were stained by 10 µg/ml Hoechst 33342 for 8 min. Finally, the images were collected by an EVOS fluorescence microscope. The information of antibodies used in this study was listed in Table S4.

### Chromatin Immunoprecipitation and ChIP-seq

#### ChIP Assay

The 1×10^7^ cells were crosslinked with 1% formaldehyde for 10 min at room temperature, then added to 125 mM Glycine to neutralize the formaldehyde. The crosslinked cells were washed three times using ice-cold PBS, subsequently scraped off and transferred to a 15ml centrifuge tube. The cell suspension was spun down for 5 min at 2000g and the supernatant was completely removed. The sediments were lysed by Nuclear Lysis Buffer to obtain chromatin extracts, that were treated with ultrasound to obtain DNA fragments with an average size of 200-500 bp. The complexes of target proteins and DNA fragments are immunoprecipitated by specific antibodies and protein A/G magnetic beads. The information of antibodies used in this study were listed in Table S4.

#### ChIP-seq and data analysis

The DNA fragments of IP and Input were used for stranded DNA library preparation. Then the library was detected by agarose electrophoresis, quantified using Qubit 2.0 and sequenced on Illumina HiSeq2500 PE150. Raw sequencing data was first filtered by Trimmomatic [50] (version 0.36), low-quality reads were discarded and the reads contaminated with adaptor sequences were trimmed. The clean reads were mapped to the reference genome of*Sus scrofa* fromGCF_000003025.6(ftp://ftp.ncbi.nlm.nih.gov/genomes/all/GCF/000/003/025/GCF_000003025.6_Sscrofa11.1/GCF_000003025.6_Sscrofa11.1_genomic.fna.gz) using STAR software (version 2.5.3a) with default parameters [51]. Then the distribution, coverage, homogeneity and chain specificity of reads were evaluated by RSeQC (version 2.6) [52]. Peaks in the ChIP-Seq datasets were called with MACS2 (version 2.1.1) [53] and associated with genes using Homer (version v4.10) [53, 54]. The distribution of call peak on the chromosomes and functional elements was performed by deepTools(version 2.4.1)[55]and ChIPseeker (version 1.5.1) [56]. Peak maps of reads across the genome are described using IGV (version 2.4.16) [57]. The motif analysis was performed using Homer (version v4.10) [54]. Gene ontology (GO) analysis and Kyoto encyclopedia of genes and genomes (KEGG) enrichment analysis for the peak related genes were both implemented by KOBAS software (version: 2.1.1) with a corrected P-value cutoff of 0.05 to judge statistically significant enrichment [58]. Signal tracks of features (H3K9me2 and H3K9me3) were calculated by using Deeptools with parameter “ComputeMatrix”. We remove the noise by subtracting the INPUT matrix from the IP matrix and showed the features at peaks regions and up- and downstream regions within ±2kb.

### Digital RNA-seq

#### RNA extraction, library preparation and sequencing

Total RNAs were extracted from samples using TRIzol (Invitrogen) following the methods [59]. DNA digestion was carried out after RNA extraction by DNase I. RNA quality was determined by examining A260/A280 with NanodropTM One spectrophotometer (Thermo Fisher Scientific Inc). RNA Integrity was confirmed by 1.5% agarose gel electrophoresis. Qualified RNAs were finally quantified by Qubit3.0 with QubitTM RNA Broad Range Assay kit (Life Technologies).

2 μg total RNAs were used for stranded RNA sequencing library preparation using KC-DigitalTM Stranded mRNA Library Prep Kit for Illumina® (Catalog NO. DR08502, Wuhan Seqhealth Co., Ltd. China) following the manufacturer’s instruction. The kit eliminates duplication bias in PCR and sequencing steps, by using unique molecular identifier (UMI) of 8 random bases to label the pre-amplified cDNA molecules. The library products corresponding to 200-500 bps were enriched, quantified and finally sequenced on Hiseq X 10 sequencer (Illumina).

#### RNA-Seq data analysis

Raw sequencing data was first filtered by Trimmomatic (version 0.36) [50], low-quality reads were discarded and the reads contaminated with adaptor sequences were trimmed. Clean Reads were further treated with in-house scripts to eliminate duplication bias introduced in library preparation and sequencing. Briefly, clean reads were first clustered according to the UMI sequences, in which reads with the same UMI sequence were grouped into the same cluster, resulting in 65,536 clusters. Reads in the same cluster were compared to each other by pairwise alignment, and then reads with sequence identity over 95% were extracted to a new sub-cluster. After all sub-clusters were generated, multiple sequence alignment was performed to get one consensus sequence for each sub-cluster. After these steps, any errors and biases introduced by PCR amplification or sequencing were eliminated [60, 61].

The de-duplicated consensus sequences were used for standard RNA-seq analysis. They were mapped to the reference genome of *Sus scrofa* from GCF_000003025.6(ftp://ftp.ncbi.nlm.nih.gov/genomes/all/GCF/000/003/025/GCF_000003025.6_Sscrofa11.1/GCF_000003025.6_Sscrofa11.1_genomic.fna.gz) using STAR software (version 2.5.3a) [51] with default parameters. Reads mapped to the exon regions of each gene were counted by feature Counts (Version 1.5.1) [62] and then FPKMs were calculated. Genes differentially expressed between groups were identified using the edgeR package (version 3.12.1) [63]. An FDR corrected p-value?cutoff of 0.05 and Fold-change cutoff of 2 were used to judge the statistical significance of gene expression differences. Gene ontology (GO) analysis and Kyoto encyclopedia of genes and genomes (KEGG) enrichment analysis for differentially expressed genes were both implemented by KOBAS software (version: 2.1.1) [58] with a corrected P-value cutoff of 0.05 to judge statistically significant enrichment. We clustered and showed our data through hierarchical clustering of R package “pheatmap” with parameter “cutree_rows” based on cluster results. Hierarchical clustering was performed with a distance matrix between two parts. Distance measure used in clustering rows by calculating Pearson correlation.

#### ATAC-seq

5 × 10^4^ cells were treated with cell lysis buffer and nucleus was collected by density gradient centrifuging for 30min at 120000g in 60% Percoll and 2.5M sucrose solution. Transposition and high-throughput DNA sequencing library were carried out by TruePrep DNA Library Prep Kit V2 for Illumina kit (Catalog NO. TD501, Vazyme). The library was sequenced on Novaseq 6000 sequencer (Illumina).

Raw sequencing data was first filtered by Trimmomatic (version 0.36). Clean Reads were further treated with FastUniq (version 1.1) to eliminate duplication. Reads were mapped to the reference genome of Sus scrofa from GCF_000003025.6 using bowtie2 software (version 2.2.6) with default parameters. The MACS2 software (version 2.1.1) was used for peak calling. The Homer (version 4.10) was used for motifs analysis. We got OS binding sites from ChIP-seq of O/S in this study. Then, genome signal tracks matrix of ATAC-seq nearby O/S sites were calculated by computeMatrix of deeptools. Signal of ATAC-seq nearby O/S sites were visualized by ggplot2.

### IP-MS

#### Immunoprecipitation

1×10^7^ cells were digested by TrypLE™ Select and then transferred immediately to a 1.5 ml tube on ice. The cell suspension was then centrifuged at 5000g for 3 min and supernatant was discarded. The cell sediment was lysed by IP buffer (Beyotime, P0013) for 30 min on ice and then centrifuged at 12000g for 10 min and supernatant was collected. The supernatant was incubated with anti-FLAG magnetic beads (Sigma, F2426) overnight at 4°C, washed five times with IP buffer, added to 5× SDS-PAGE loading buffer (GENSHARE G, JC-PE007), and heated at 100 °C for 5 min. Immunoprecipitates were subjected to SDS-PAGE and probed with indicated antibodies.

### Mass Spectrometry

#### In-gel Digestion

For in-gel tryptic digestion, gel pieces were destained in 50 mM NH4HCO3 in 50% acetonitrile (v/v) until clear. Gel pieces were dehydrated with 100 μl of 100% acetonitrile for 5 min, the liquid was removed, and the gel pieces rehydrated in 10 mM dithiothreitol and incubated at 56 °C for 60 min. Gel pieces were again dehydrated in 100% acetonitrile, the liquid was removed and gel pieces were rehydrated with 55 mM iodoacetamide. Samples were incubated at room temperature in the dark for 45 min. Gel pieces were washed with 50 mM NH4HCO3 and dehydrated with 100% acetonitrile. Gel pieces were rehydrated with 10 ng/μl trypsin resuspended in 50 mM NH4HCO3 on ice for 1 h. Excess liquid was removed and gel pieces were digested with trypsin at 37 °C overnight. Peptides were extracted with 50% acetonitrile/5% formic acid, followed by 100% acetonitrile. Peptides were dried to completion and resuspended in 2% acetonitrile/0.1% formic acid.

#### LC-MS/MS Analysis

The tryptic peptides were dissolved in 0.1% formic acid (solvent A), directly loaded onto a home-made reversed-phase analytical column (15 cm length, 75 μm i.d.). The gradient was comprised of an increase from 6% to 23% solvent B (0.1% formic acid in 98% acetonitrile) over 16 min, 23% to 35% in 8 min and climbing to 80% in 3 min then holding at 80% for the last 3 min, all at a constant flow rate of 400 nl/min on an EASY-nLC 1000 UPLC system.

The peptides were subjected to NSI source followed by tandem mass spectrometry (MS/MS) in Q ExactiveTM Plus (Thermo) coupled online to the UPLC. The electrospray voltage applied was 2.0 kV. The m/z scan range was 350 to 1800 for full scan, and intact peptides were detected in the Orbitrap at a resolution of 70,000. Peptides were then selected for MS/MS using NCE setting as 28 and the fragments were detected in the Orbitrap at a resolution of 17,500. A data-dependent procedure that alternated between one MS scan followed by 20 MS/MS scans with 15.0s dynamic exclusion. Automatic gain control (AGC) was set at 5E4.

### Data Processing

The resulting MS/MS data were processed using Proteome Discoverer 1.3. Tandem mass spectra were searched against UniPort database. Trypsin/P (or other enzymes if any) was specified as cleavage enzyme allowing up to 2 missing cleavages. The mass error was set to 10 ppm for precursor ions and 0.02 Da for-fragment ions. Carbamidomethyl on Cys were specified as fixed modification and oxidation on Met was specified as variable modification. Peptide confidence was set at high, and peptide ion score was set > 20.

### Luciferase assay for enhancer activity

All Luciferase vectors were derived from PGL3-basic vector (Promega, E1751) by Seamless Cloning. Luciferase activity was measured using dual-luciferase detection kit (Beyotime, RG027) as described in the manufacturer’s procedure.

### Statistical analysis

All the experiments had three independent biological replicates, except RNA-seq which were replicated twice. The ChIP samples were collected from three independent biological experiments, then mixed for ChIP-seq. Statistical significance was accepted at P < 0.05 and calculated using one-way ANOVA in Excel 2016.

## Acknowledgments

We thank Dr. Yang Fan’s helpful review and comments on the manuscript. We thank seqhealth technology co., LTD supplies technical support.

## Author contributions

Zhenshuo Zhu and Jinlian Hua designed the study and wrote the manuscript. Zhenshuo Zhu, Xiaolong Wu, Juqing Zhang, Shuai Yu, Qiaoyan Shen, Zhe Zhou, Qin Pan, Wei Yue, Dezhe Qin, Ying Zhang, Wenxu Zhao, Rui Zhang performed the experiments. Zhenshuo Zhu, Qun Li, Sha Peng, Na Li, Shiqiang Zhang, Anmin Lei, Yiliang Miao, Zhonghua Liu, Huayan Wang, Mingzhi Liao, Jinlian Hua analyzed the data.

## Data availability

ChIP-seq and RNA-Seq data have been uploaded to National Center for Biotechnology Information Sequence Read Archive (SRA) database under the accession number PRJNA633419. The mass spectrometry proteomics data have been deposited to the ProteomeXchange Consortium (http://proteomecentral.proteomexchange.org) via the iProX partner repository [64] with the dataset identifier PXD016937.

