## supplemental figures and tables for "Histone demethylase complexes KDM3A and KDM3B cooperate with OCT4/SOX2 to construct pluripotency gene regulatory network"

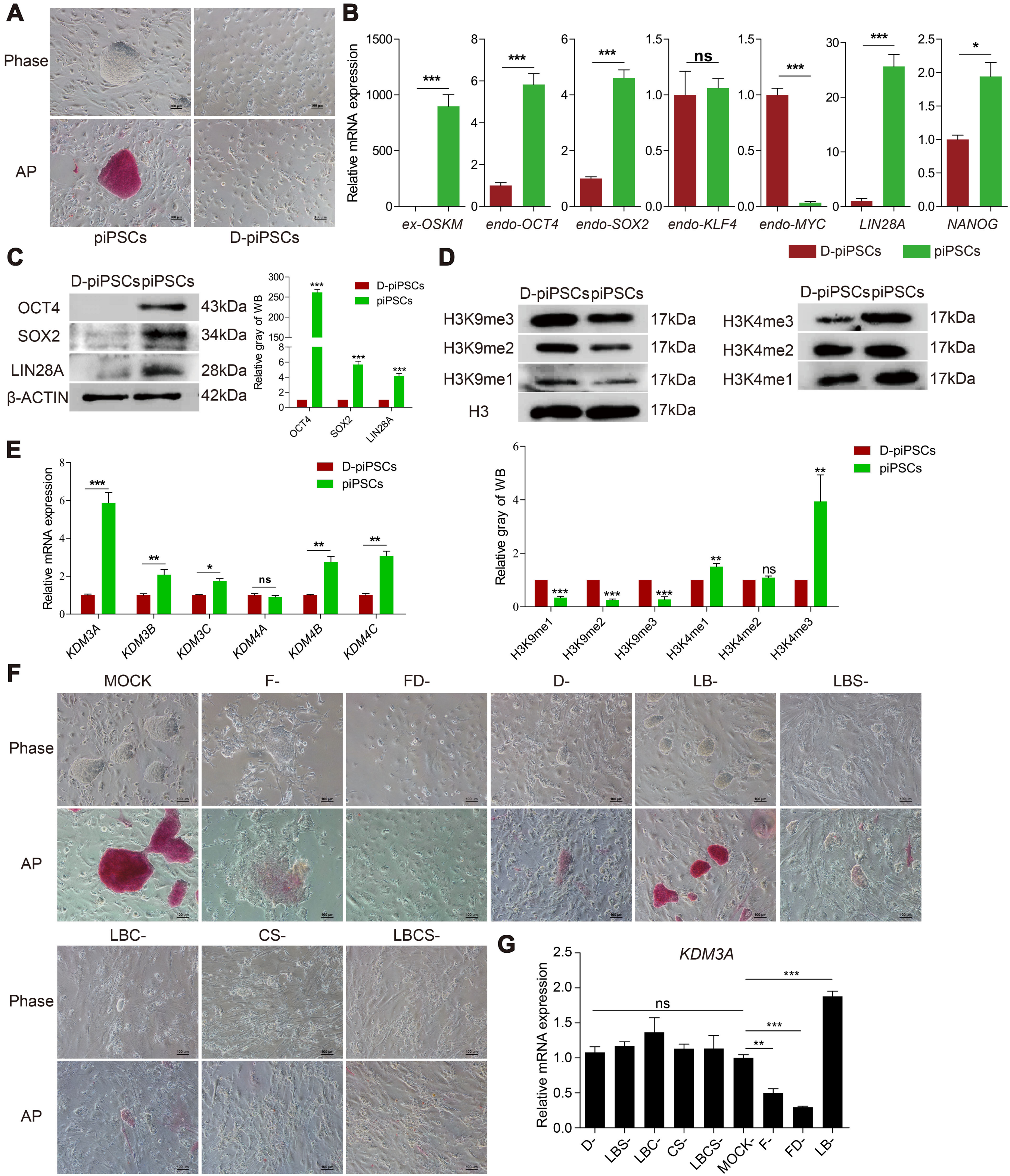
A
Phase
AP
piPSCs
D-piPSCs
B
Relative mRNA expression
ex-OSKM
endo-OCT4
endo-SOX2
endo-KLF4
endo-MYC
LIN28A
NANOG
ns
D-piPSCs
piPSCs
C
D-piPSCs piPSCs
OCT4
SOX2
LIN28A
β-ACTIN
43kDa
34kDa
28kDa
42kDa
Relative gray of WB
D
H3K9me3
H3K9me2
H3K9me1
H3
H3K4me3
H3K4me2
H3K4me1
17kDa
E
KDM3A
KDM3B
KDM3C
KDM4A
KDM4B
KDM4C
H3K9me1
H3K9me2
H3K9me3
H3K4me1
H3K4me2
H3K4me3
F
MOCK
F-
FD-
D-
LB-
LBS-
LBC-
CS-
LBCS-
G
KDM3A
D-
LBS-
LBC-
CS-
LBCS-
MOCK-
F-
FD-
LB-

### Supplementary figures

**Fig. S1. The loss of pluripotency was accompanied by the increase of H3K9 methylation and the decrease of corresponding demethylase. (A)** Representative image of AP staining of piPSCs and D-piPSCs (differentiated piPSCs). The D-piPS represents piPSCs after 5 days of grown under differentiation medium. The experiments were performed three times. The scale bar represents 100 μm. **(B)** RT–qPCR analysis of the exogenous reprogramming factors and endogenous pluripotent genes in the piPS and D-piPS. The relative expression levels were normalized to β-actin. Data represent the mean ± s.d; n = 3 independent experiments. **(C)** Representative Western-Blot of OCT4, SOX2, LIN28A after 5 days of culture in the indicated conditions. The quantitative analysis is shown by bar graph. Data represent the mean ± s.d; n = 3 independent experiments. **(D)** Representative Western-Blot of H3K9me1/2/3 and H3K4me1/2/3 after 5 days of culture in the indicated conditions. The quantitative analysis is shown by bar graph. Data represent the mean ± s.d; n = 3 independent experiments. **(E)** RT–qPCR analysis of *KDM3A/3B/3C* and *KDM4A/B/C* in the piPS and D-piPS. The relative expression levels were normalized to β-actin. Data represent the mean ± s.d.; n = 3 independent experiments. **(F)** piPSCs were cultured in medium with or without LIF, b-FGF, Chir99021, SB431542, feeder and DOX and detected by AP staining. MOCK: piPSCs grown under normal condition as a blank control; F-: feeder free; FD-: feeder free and absence of DOX; D-: absence of DOX; LB-: absence of LIF and bFGF; LBS-: absence of LIF, bFGF and SB431542; LBC-: absence of LIF, bFGF and Chir99021; CS-: absence of Chir99021 and SB431542; LBCS-: absence of LIF, bFGF, Chir99021 and SB431542. **(G)** RT–qPCR analysis of *KDM3A* in the indicated conditions. The relative expression levels were normalized to β-actin. Data represent the mean ± s.d.; n = 3 independent experiments.

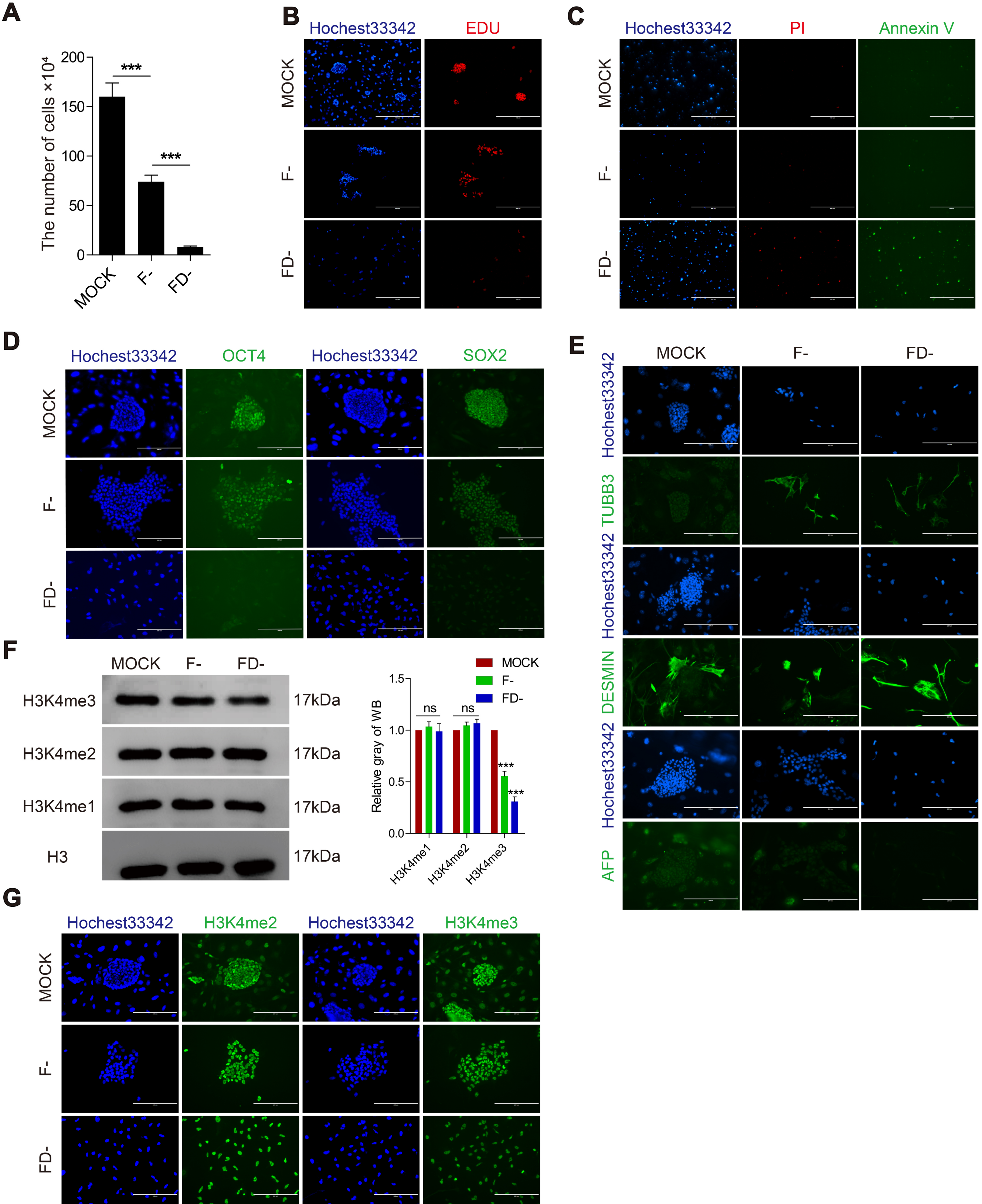
A
The number of cells ×10⁴
200
150
100
50
0
***
***
MOCK
F-
FD-
B
Hochest33342
EDU
MOCK
F-
FD-
C
Hochest33342
PI
Annexin V
MOCK
F-
FD-
D
Hochest33342
OCT4
Hochest33342
SOX2
MOCK
F-
FD-
E
MOCK
F-
FD-
Hochest33342
TUBB3
Hochest33342
DESMIN
Hochest33342
AFP
F
MOCK
F-
FD-
H3K4me3
H3K4me2
H3K4me1
H3
17kDa
17kDa
17kDa
17kDa
Relative gray of WB
1.5
1.0
0.5
0.0
ns
ns
***
***
MOCK
F-
FD-
H3K4me1
H3K4me2
H3K4me3
G
Hochest33342
H3K4me2
Hochest33342
H3K4me3
MOCK
F-
FD-

**Fig. S2. The phenotype of pips showed gradual differentiation, decreased proliferation rate and death with F- and FD- treatments. (A)** The number of total cells in MOCK, F- and FD- conditions on day 5. Data represent the mean ± s.d.; n = 3 independent experiments. **(B)** Representative images of cells treated with F- and FD- and exposed to EDU in order to evaluate proliferation. n = 3 independent experiments. **(C)** Representative images of cells stained with PI and annexin V after 5 days of culture in MOCK, F- and FD- conditions. n = 3 independent experiments. **(D)** Immunofluorescence analysis of OCT4 and SOX2 in the indicated conditions. The nuclei were DAPI stained. The scale bar represents 200 μm. The experiments were performed three times. **(E)** Immunofluorescence analysis of TUBB3, Desmin and AFP in the indicated conditions. The nuclei were DAPI stained. The scale bar represents 200 μm. The experiments were performed three times. **(F)** Representative Western-Blot of H3K4me1/2/3 after 5 days of culture in the indicated conditions. The quantitative analysis is shown by bar graph. Data represent the mean ± s.d.; n = 3 independent experiments. **(G)** Immunofluorescence analysis of H3K4me1/2/3 in the indicated conditions. The nuclei were DAPI stained. The scale bar represents 200 μm. The experiments were performed three times.

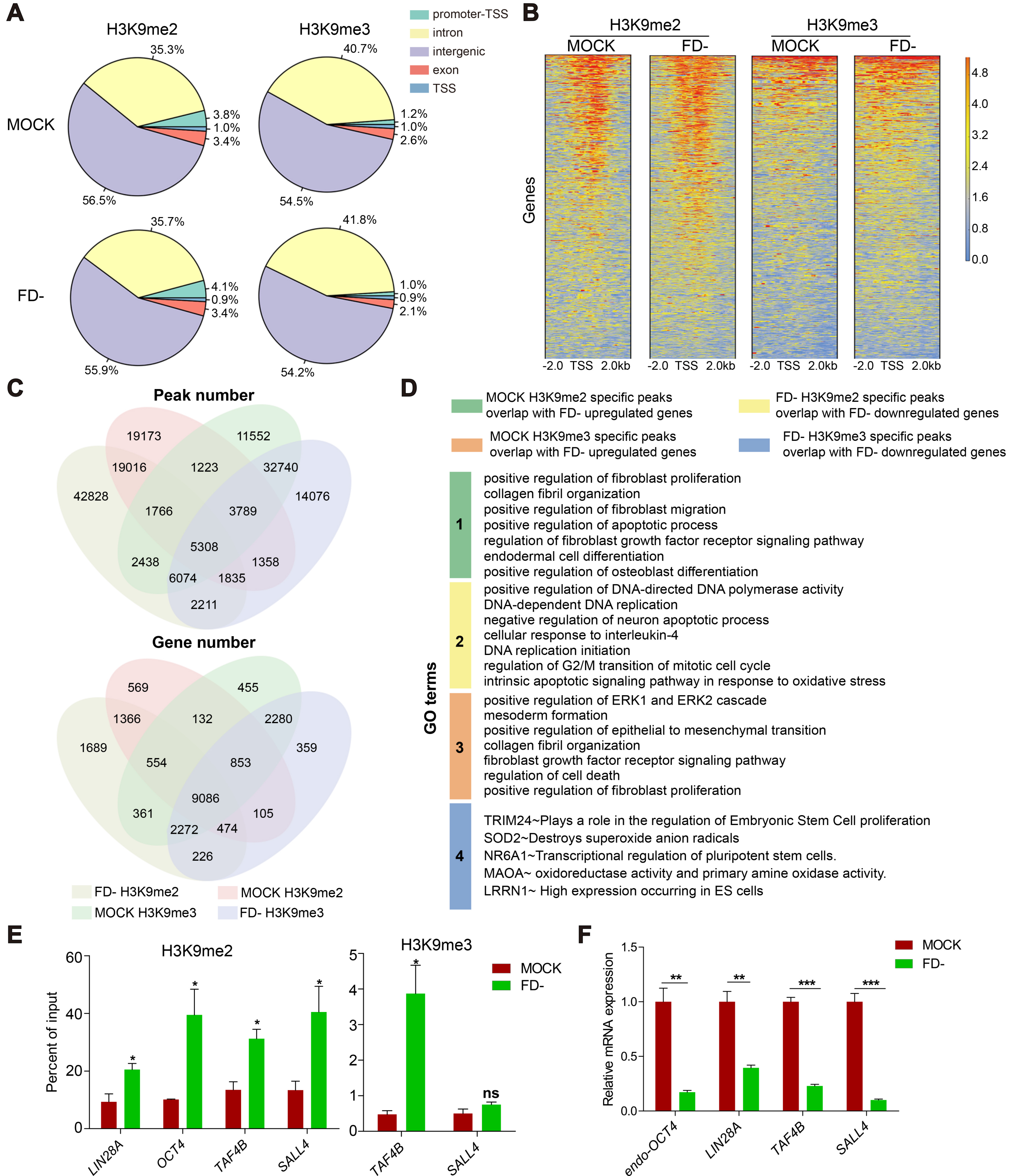
A
H3K9me2
H3K9me3
promoter-TSS
intron
intergenic
exon
TSS
MOCK
35.3%
3.8%
1.0%
3.4%
56.5%
40.7%
1.2%
1.0%
2.6%
54.5%
FD-
35.7%
4.1%
0.9%
3.4%
55.9%
41.8%
1.0%
0.9%
2.1%
54.2%
B
H3K9me2
H3K9me3
MOCK
FD-
MOCK
FD-
Genes
-2.0 TSS 2.0kb
4.8
4.0
3.2
2.4
1.6
0.8
0.0
C
Peak number
19173
11552
19016
1223
32740
42828
14076
1766
3789
5308
2438
1358
6074
1835
2211
Gene number
569
455
1366
132
2280
1689
359
554
853
9086
361
105
2272
474
226
FD- H3K9me2
MOCK H3K9me2
MOCK H3K9me3
FD- H3K9me3
D
MOCK H3K9me2 specific peaks overlap with FD- upregulated genes
FD- H3K9me2 specific peaks overlap with FD- downregulated genes
MOCK H3K9me3 specific peaks overlap with FD- upregulated genes
FD- H3K9me3 specific peaks overlap with FD- downregulated genes
GO terms
1
positive regulation of fibroblast proliferation
collagen fibril organization
positive regulation of fibroblast migration
positive regulation of apoptotic process
regulation of fibroblast growth factor receptor signaling pathway
endodermal cell differentiation
positive regulation of osteoblast differentiation
2
positive regulation of DNA-directed DNA polymerase activity
DNA-dependent DNA replication
negative regulation of neuron apoptotic process
cellular response to interleukin-4
DNA replication initiation
regulation of G2/M transition of mitotic cell cycle
intrinsic apoptotic signaling pathway in response to oxidative stress
3
positive regulation of ERK1 and ERK2 cascade
mesoderm formation
positive regulation of epithelial to mesenchymal transition
collagen fibril organization
fibroblast growth factor receptor signaling pathway
regulation of cell death
positive regulation of fibroblast proliferation
4
TRIM24~Plays a role in the regulation of Embryonic Stem Cell proliferation
SOD2~Destroys superoxide anion radicals
NR6A1~Transcriptional regulation of pluripotent stem cells.
MAOA~ oxidoreductase activity and primary amine oxidase activity.
LRRN1~ High expression occurring in ES cells
E
H3K9me2
Percent of input
60
40
20
0
LIN28A
OCT4
TAF4B
SALL4
H3K9me3
5
4
3
2
1
0
TAF4B
SALL4
ns
MOCK
FD-
F
Relative mRNA expression
1.5
1.0
0.5
0.0
**
**
***
***
endo-OCT4
LIN28A
TAF4B
SALL4
MOCK
FD-

**Fig. S3. The dynamic distribution of H3K9me2/3 changes between MOCK and FD- treatments.** **(A)** Fraction of H3K9me2/3 binding sites within various regions of the genome. **(B)** Distribution of H3K9me2/3 ChIP-seq signal on both sides of transcription start sites (TSS). **(C)** Venn diagrams showing the genome-wide overlap of the H3K9me2/3 binding sites and its associated genes between MOCK and FD- conditions. **(D)** Combined analysis of H3K9me2/3 ChIP-seq and RNA-seq. The downregulated genes of FD- overlap with the specific H3K9me2/3 peaks of FD-. The upregulated genes of FD- overlap with the specific H3K9me2/3 peaks of MOCK. Gene ontology enrichment for each cluster is presented on the right. **(E)** ChIP–qPCR analysis of *OCT4, LIN28A, SALL4* and *TAF4B* in the indicated conditions. The bold black line in Fig. 3D indicates the detection regions of ChIP-qPCR. Data represent the mean ± s.d.; n = 3 independent experiments. **(F)** RT–qPCR analysis of *endo-OCT4, LIN28A, SALL4* and *TAF4B* in the indicated conditions. The relative expression levels were normalized to β-actin. Data represent the mean ± s.d.; n = 3 independent experiments.

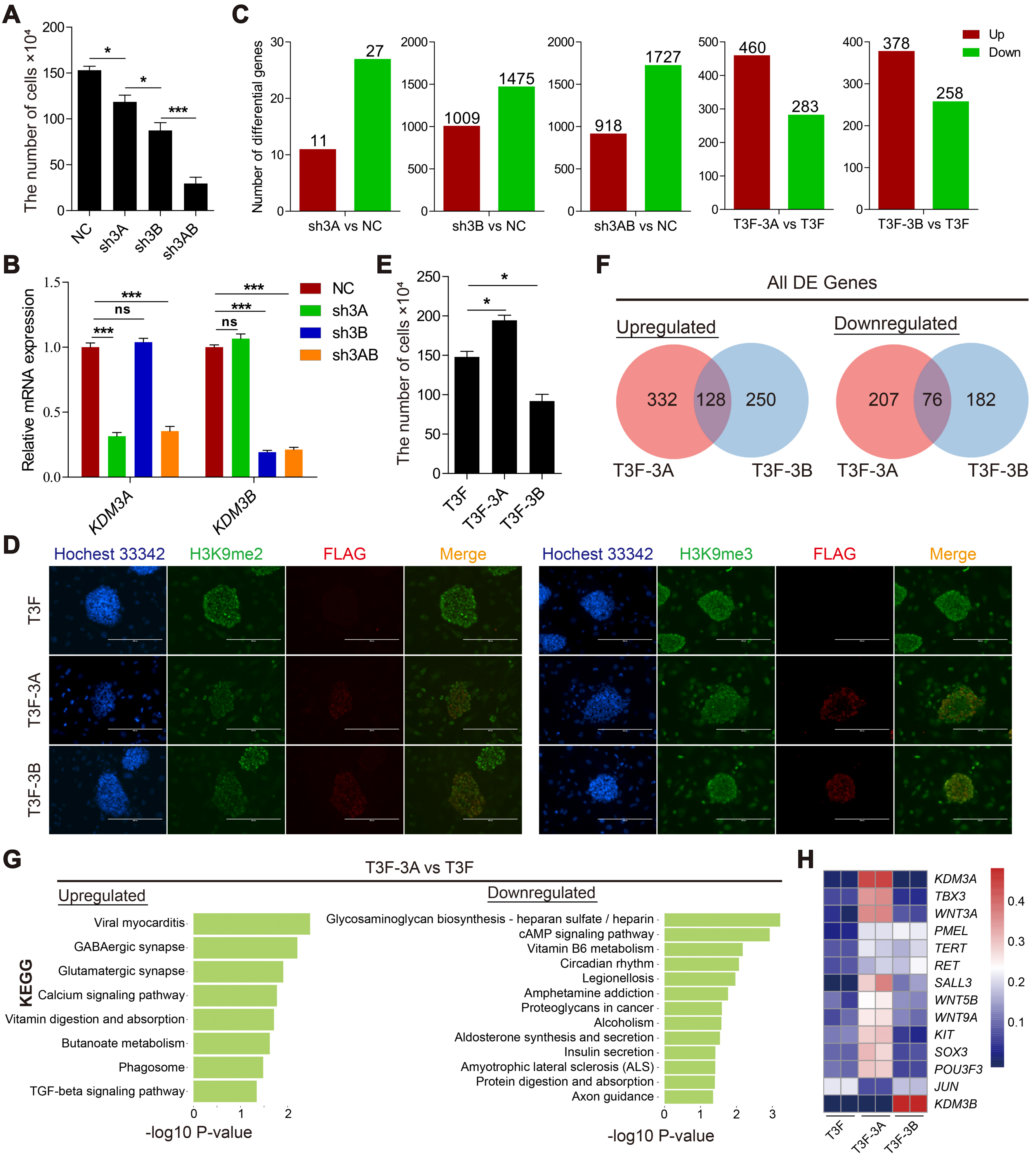
A
The number of cells ×10⁴
NC
sh3A
sh3B
sh3AB
C
Number of differential genes
11
27
sh3A vs NC
1009
1475
sh3B vs NC
918
1727
sh3AB vs NC
460
283
T3F-3A vs T3F
378
258
T3F-3B vs T3F
Up
Down
B
Relative mRNA expression
NC
sh3A
sh3B
sh3AB
KDM3A
KDM3B
E
The number of cells ×10⁴
T3F
T3F-3A
T3F-3B
F
All DE Genes
Upregulated
Downregulated
332
128
250
207
76
182
T3F-3A
T3F-3B
D
Hochest 33342
H3K9me2
FLAG
Merge
H3K9me3
G
T3F-3A vs T3F
KEGG
Viral myocarditis
GABAergic synapse
Glutamatergic synapse
Calcium signaling pathway
Vitamin digestion and absorption
Butanoate metabolism
Phagosome
TGF-beta signaling pathway
Glycosaminoglycan biosynthesis - heparan sulfate / heparin
cAMP signaling pathway
Vitamin B6 metabolism
Circadian rhythm
Legionellosis
Amphetamine addiction
Proteoglycans in cancer
Alcoholism
Aldosterone synthesis and secretion
Insulin secretion
Amyotrophic lateral sclerosis (ALS)
Protein digestion and absorption
Axon guidance
-log10 P-value
H
KDM3A
TBX3
WNT3A
PMEL
TERT
RET
SALL3
WNT5B
WNT9A
KIT
SOX3
POU3F3
JUN
KDM3B

**Fig. S4. KDM3A/B promotes transcriptional output by reducing H3K9me2/3. (A)** The number of total cells of the NC, sh3A, sh3B and sh3AB cell lines after 5 days of clonal growth. Data represent the mean ± s.d.; n = 3 independent experiments. **(B)** RT–qPCR analysis of the *KDM3A* and *KDM3B* in the NC, sh3A, sh3B and sh3AB cell lines. The relative expression levels were normalized to β-actin. Data represent the mean ± s.d; n = 3 independent experiments. **(C)** Bar graph of the number of differentially expressed genes. **(D)** Immunofluorescence analysis of H3K9me2/3 in the indicated conditions. The nuclei were DAPI stained. The scale bar represents 200 μm. The experiments were performed three times. **(E)** The number of total cells of the T3F, T3F-3A and T3F-3B cell lines after 5 days of clonal growth. Data represent the mean ± s.d.; n = 3 independent experiments. **(F)** Venn diagrams showing the overlap of up- or downregulated differentially expressed (DE) transcripts between T3F-3A and T3F-3B cell lines on day 5 determined by RNA-Seq. **(G)** KEGG enrichment of up- or downregulated differentially expressed (DE) genes in T3F-3B cell line. **(H)** Heatmap showing changes in the expression of some differentially expressed genes in the T3F, T3F-3A and T3F-3B cell lines obtained from RNA-Seq analysis. Data were normalized by dividing the FPKM of each gene by the sum.

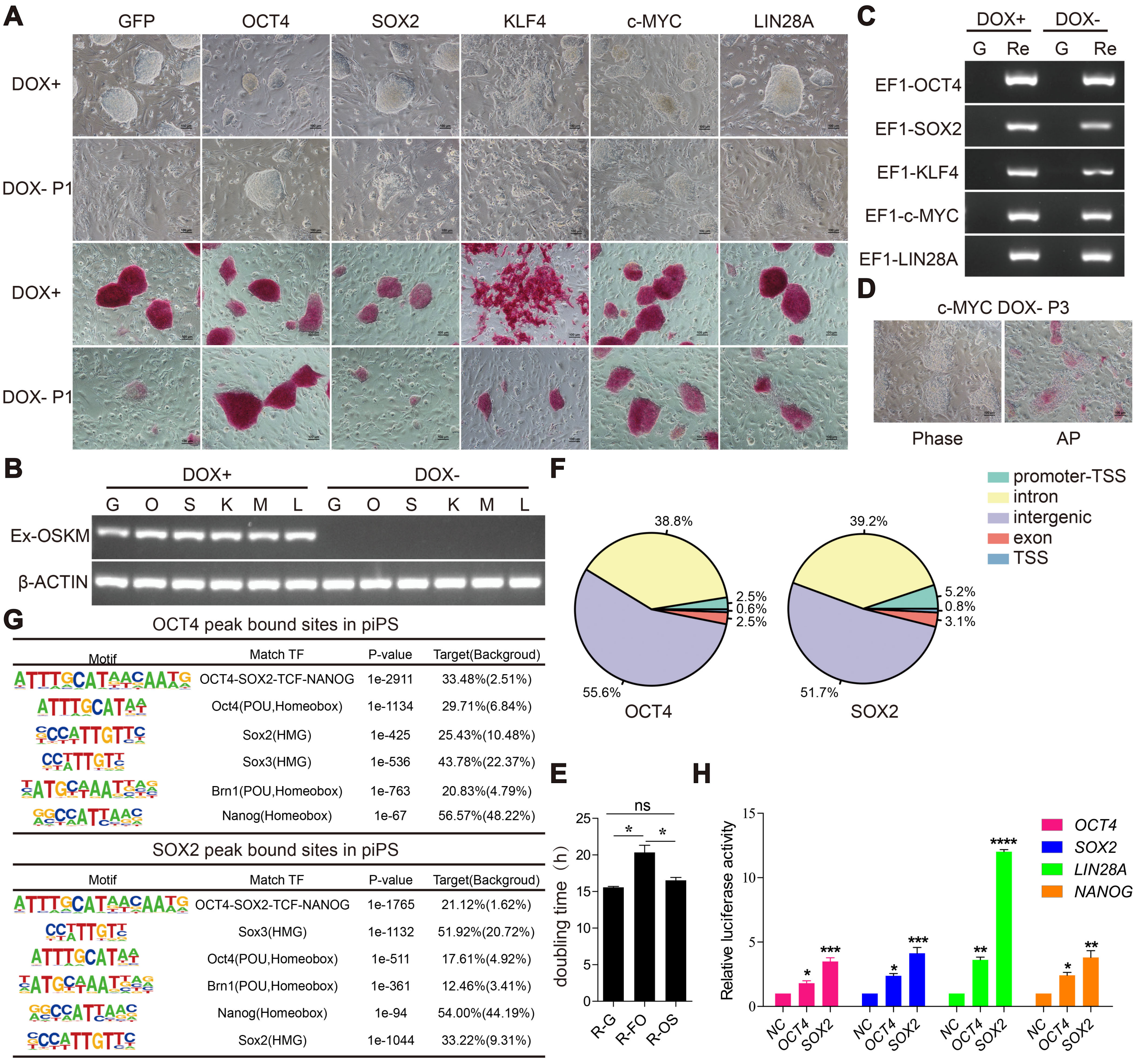
A
GFP
OCT4
SOX2
KLF4
c-MYC
LIN28A
DOX+
DOX- P1
DOX+
DOX- P1
C
DOX+
DOX-
G
Re
G
Re
EF1-OCT4
EF1-SOX2
EF1-KLF4
EF1-c-MYC
EF1-LIN28A
D
c-MYC DOX- P3
Phase
AP
B
DOX+
DOX-
G O S K M L G O S K M L
Ex-OSKM
β-ACTIN
F
promoter-TSS
intron
intergenic
exon
TSS
38.8%
2.5%
0.6%
2.5%
55.6%
OCT4
39.2%
5.2%
0.8%
3.1%
51.7%
SOX2
G
OCT4 peak bound sites in piPS
Motif | Match TF | P-value | Target(Backgroud)
OCT4-SOX2-TCF-NANOG | 1e-2911 | 33.48%(2.51%)
Oct4(POU,Homeobox) | 1e-1134 | 29.71%(6.84%)
Sox2(HMG) | 1e-425 | 25.43%(10.48%)
Sox3(HMG) | 1e-536 | 43.78%(22.37%)
Brn1(POU,Homeobox) | 1e-763 | 20.83%(4.79%)
Nanog(Homeobox) | 1e-67 | 56.57%(48.22%)
SOX2 peak bound sites in piPS
Motif | Match TF | P-value | Target(Backgroud)
OCT4-SOX2-TCF-NANOG | 1e-1765 | 21.12%(1.62%)
Sox3(HMG) | 1e-1132 | 51.92%(20.72%)
Oct4(POU,Homeobox) | 1e-511 | 17.61%(4.92%)
Brn1(POU,Homeobox) | 1e-361 | 12.46%(3.41%)
Nanog(Homeobox) | 1e-94 | 54.00%(44.19%)
Sox2(HMG) | 1e-1044 | 33.22%(9.31%)
E
ns
*
*
doubling time (h)
0
5
10
15
20
25
R-G
R-FO
R-OS
H
Relative luciferase activity
0
5
10
15
OCT4
SOX2
LIN28A
NANOG
*
***
*
***
**
****
*
**
NC OCT4 SOX2
NC OCT4 SOX2
NC OCT4 SOX2
NC OCT4 SOX2

**Fig. S5. The regulatory model among key pluripotent factors in piPSCs. (A)** Representative image of AP stained colonies after 5 days of clonal growth of indicated cell lines in the DOX+ and DOX- P1 (absence of Dox, passage 1) treatments. EF1-GFP(GFP), EF1-OCT4(OCT4), EF1-SOX2 (SOX2), EF1-KLF4 (KLF4) and EF1-c-MYC(c-MYC) represents the corresponding gene that is stably overexpressed in piPSCs. GFP was used as a negative control. The experiments were performed three times. The scale bar represents 100 μm. **(B)** RT-PCR analysis of insert gene expression in the indicated cell lines. **(C)** RT–PCR analysis of the exogenous reprogramming factors in the indicated conditions. **(D)** Representative image of AP stained colonies after 5 days of clonal growth of c-MYC cell lines in the DOX- P3 (absence of Dox, passage 3) treatments. The scale bar represents 100 μm. **(E)** The doubling time of R-G, R-FO and R-OS cell lines. Data are from three biological replicates and are shown as the mean ± s.d. **(F)** Fraction of OCT4 and SOX2 binding sites within various regions of the genome. **(G)** The motifs of OCT4 and SOX2 were identified in piPSCs. **(H)** Enhancer-luciferase assays for the OCT4 and SOX2 binding regions of OCT4, SOX2, LIN28A and NANOG. NC, Empty Vector. Data are from three biological replicates and are shown as the mean mean ± s.d.

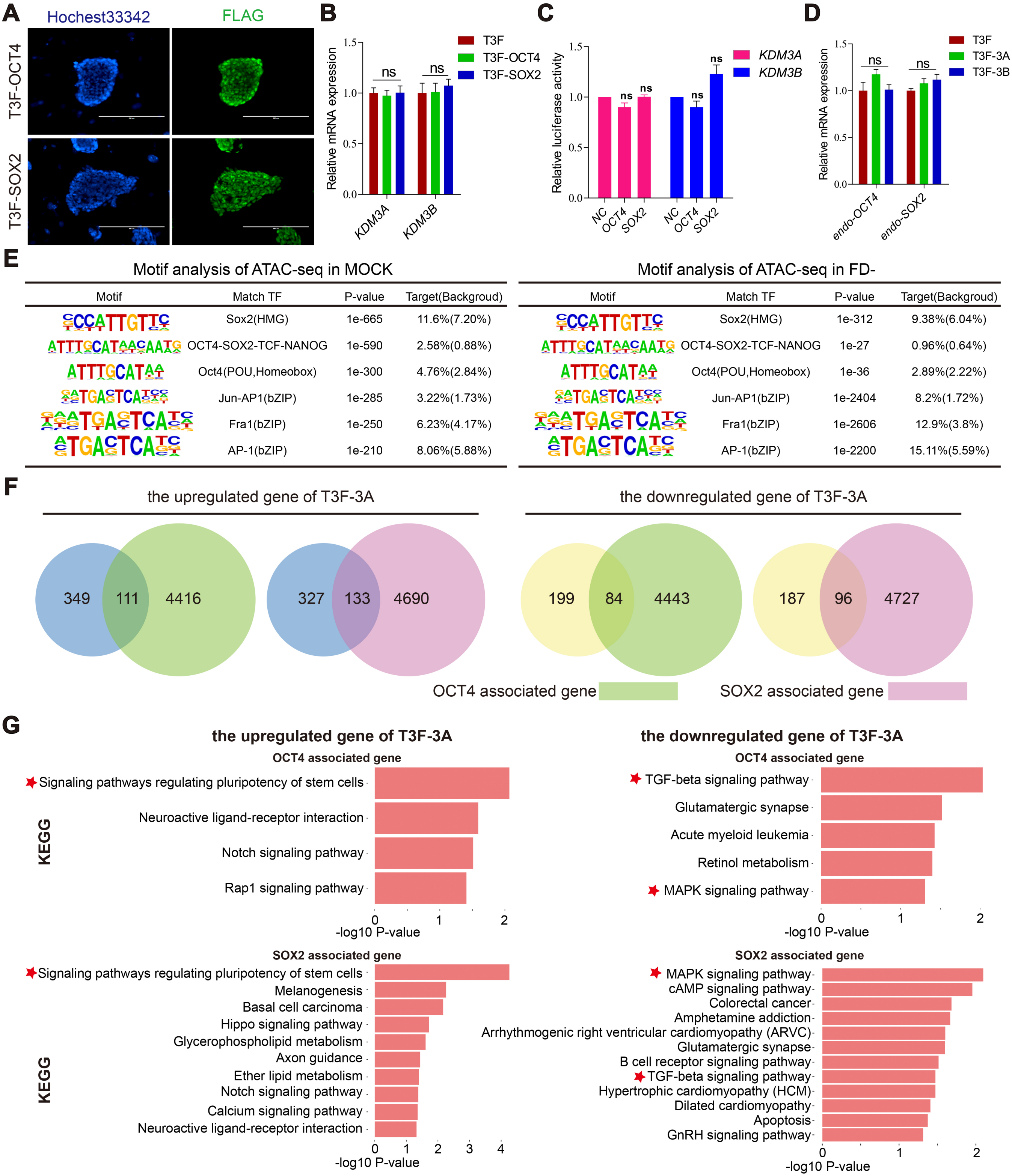
A
Hochest33342
FLAG
T3F-OCT4
T3F-SOX2
B
Relative mRNA expression
T3F
T3F-OCT4
T3F-SOX2
ns
KDM3A
KDM3B
C
Relative luciferase activity
KDM3A
KDM3B
NC
OCT4
SOX2
D
Relative mRNA expression
T3F
T3F-3A
T3F-3B
endo-OCT4
endo-SOX2
E
Motif analysis of ATAC-seq in MOCK
Motif
Match TF
P-value
Target(Backgroud)
Sox2(HMG) 1e-665 11.6%(7.20%)
OCT4-SOX2-TCF-NANOG 1e-590 2.58%(0.88%)
Oct4(POU,Homeobox) 1e-300 4.76%(2.84%)
Jun-AP1(bZIP) 1e-285 3.22%(1.73%)
Fra1(bZIP) 1e-250 6.23%(4.17%)
AP-1(bZIP) 1e-210 8.06%(5.88%)
Motif analysis of ATAC-seq in FD-
Sox2(HMG) 1e-312 9.38%(6.04%)
OCT4-SOX2-TCF-NANOG 1e-27 0.96%(0.64%)
Oct4(POU,Homeobox) 1e-36 2.89%(2.22%)
Jun-AP1(bZIP) 1e-2404 8.2%(1.72%)
Fra1(bZIP) 1e-2606 12.9%(3.8%)
AP-1(bZIP) 1e-2200 15.11%(5.59%)
F
the upregulated gene of T3F-3A
349 111 4416
327 133 4690
the downregulated gene of T3F-3A
199 84 4443
187 96 4727
OCT4 associated gene
SOX2 associated gene
G
the upregulated gene of T3F-3A
OCT4 associated gene
KEGG
Signaling pathways regulating pluripotency of stem cells
Neuroactive ligand-receptor interaction
Notch signaling pathway
Rap1 signaling pathway
-log10 P-value
SOX2 associated gene
Signaling pathways regulating pluripotency of stem cells
Melanogenesis
Basal cell carcinoma
Hippo signaling pathway
Glycerophospholipid metabolism
Axon guidance
Ether lipid metabolism
Notch signaling pathway
Calcium signaling pathway
Neuroactive ligand-receptor interaction
-log10 P-value
the downregulated gene of T3F-3A
OCT4 associated gene
TGF-beta signaling pathway
Glutamatergic synapse
Acute myeloid leukemia
Retinol metabolism
MAPK signaling pathway
-log10 P-value
SOX2 associated gene
MAPK signaling pathway
cAMP signaling pathway
Colorectal cancer
Amphetamine addiction
Arrhythmogenic right ventricular cardiomyopathy (ARVC)
Glutamatergic synapse
B cell receptor signaling pathway
TGF-beta signaling pathway
Hypertrophic cardiomyopathy (HCM)
Dilated cardiomyopathy
Apoptosis
GnRH signaling pathway
-log10 P-value

**Fig. S6. OCT4 and SOX2 cooperate with KDM3A to promote transcription output and maintain porcine pluripotent networks. (A)** Immunofluorescence analysis of Flag in the T3F-OCT4(TetO-3×flag-OCT4) and T3F-SOX2 (TetO-3×flag-SOX2). The nuclei were DAPI stained. The scale bar represents 200 μm. **(B)** RT–qPCR analysis of the *KDM3A* and *KDM3B* genes in the T3F, T3F-OCT4 and T3F-SOX2 cell lines. The relative expression levels were normalized to β-actin. Data represent the mean ± s.d; n = 3 independent experiments. **(C)** Promoter-luciferase assays for the OCT4 and SOX2 binding regions of KDM3A and KDM3B. NC, Empty Vector. Data are from three biological replicates and are shown as the mean ± s.d. **(D)** RT–qPCR analysis of the *endo-OCT4* and *endo-SOX2* in the T3F, T3F-3A and T3F-3B cell lines. The relative expression levels were normalized to β-actin. Data represent the mean ± s.d; n = 3 independent experiments. **(E)** The motifs of ATAC-seq in MOCK and FD- conditions. Last column: observed and expected motif frequencies (in parentheses). **(F)** Intersection of OCT4 or SOX2 binding genes and the upregulated and downregulated gene of T3F-3A. The number of genes for each part is given in the Venn diagram. **(G)** KEGG enrichment of intersection of OCT4 or SOX2 binding genes and the upregulated and downregulated gene of T3F-3A. The red marks represent major pluripotent regulatory pathways.

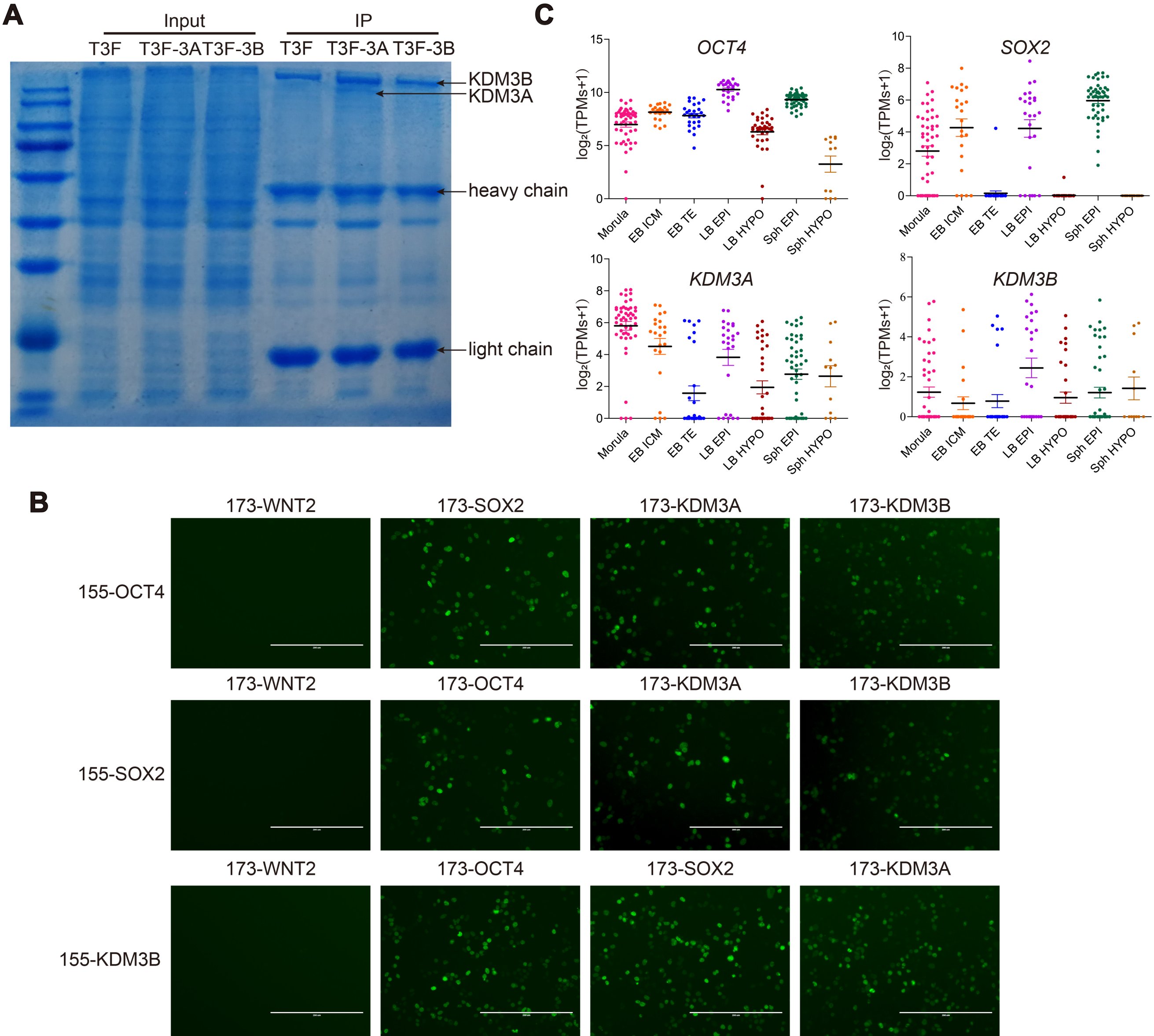
A
Input
IP
T3F
T3F-3A
T3F-3B
KDM3B
KDM3A
heavy chain
light chain
C
OCT4
SOX2
KDM3A
KDM3B
log2(TPMs+1)
Morula
EB ICM
EB TE
LB EPI
LB HYPO
Sph EPI
Sph HYPO
B
173-WNT2
173-SOX2
173-KDM3A
173-KDM3B
155-OCT4
173-OCT4
155-SOX2
155-KDM3B

**Fig. S7. OCT4 and SOX2 bind directly to the histone demethylase complex to form a transcription complex.** **(A)** KDM3A or KDM3B-interacting proteins were visualized by Coomassie brilliant blue staining. KDM3A and KDM3B were abundant in the IP; the interacting proteins were also detected and identified by mass spectrometry. **(B)** Representative image of bimolecular fluorescence complementation of OCT4, SOX2, KDM3A and KDM3B. WNT2 was used as a negative control. **(C)** Scatter plots of the expression of selected pluripotency and lineage specifier genes. The data is from (Ramos-Ibeas, 2019). EB: early blastocyst, LB: late blastocyst, Sph: spherical embryo, EPI: epiblast, HYPO: hypoblast, ICM: inner cell mass, TE: trophectoderm.

**Supplementary table 1**

**the information of vectors used in this experiment.**

| Classification for vector | Vector name | source |
|---|---|---|
| lentivirus backbones vectors | pCDH-CMV-MCS-EF1-GreenPuro | SBI, Mountain View, USA |
| | pCDH-U6-MCS-EF1-GreenPuro | our laboratory |
| | PCDH-U6-MCS-EF1-GFP-T2A- mCherry | this work |
| | pCDH-U6-shKDM3A-EF1-GreenPuro | this work |
| | pCDH-U6-shKDM3B-EF1-GreenPuro | this work |
| | PCDH-U6-shKDM3B-EF1-GFP-T2A- mCherry | this work |
| | PCDH-EF1-MCS-T2A-PURO | this work |
| | PCDH-EF1-GFP-T2A-PURO | this work |
| | PCDH-EF1-OCT4-T2A-PURO | this work |
| | PCDH-EF1-SOX2-T2A-PURO | this work |
| | PCDH-EF1-KLF4-T2A-PURO | this work |
| | PCDH-EF1-c-MYC-T2A-PURO | this work |
| | pCDH-TetO-3XFLAG-MCS-T2A-PURO | this work |
| | pCDH-TetO-3XFLAG-OCT4-T2A-PURO | this work |
| | pCDH-TetO-3XFLAG-SOX2-T2A-PURO | this work |
| | pCDH-TetO-3XFLAG-KDM3A-T2A-PURO | this work |
| | pCDH-TetO-3XFLAG-KDM3B-T2A-PURO | this work |
| Rosa26 insert vectors | Rosa26- EF1α-MCS-T2A-PURO | this work |
| | Rosa26- EF1α-GFP-T2A-PURO | this work |
| | Rosa26- EF1α-3×FLAG-OCT4-T2A-PURO | this work |
| | Rosa26- EF1α-OCT4-P2A-SOX2-T2A-PURO | this work |
| Luciferase vectors | PGL3-basic vector | Promega, E1751 |
| | PGL3-OCT4 | this work |
| | PGL3-SOX2 | this work |
| | PGL3-LIN28A | this work |
| | PGL3-NANOG | this work |
| | PGL3-KDM3A | this work |
| | PGL3-KDM3B | this work |
| BiFC vectors | pBiFC-VC155 | addgene22011 |
| | pBiFC-VN173 | addgene22010 |
| | pBiFC-VC155-OCT4 | this work |
| | pBiFC-VN173-OCT4 | this work |
| | pBiFC-VC155-SOX2 | this work |
| | pBiFC-VN173-SOX2 | this work |
| | pBiFC-VC155-KDM3B | this work |
| | pBiFC-VN173-KDM3B | this work |
| | pBiFC-VC155-KDM3A | this work |
| | pBiFC-VN173-Wnt2 | this work |

**Supplementary table 2**

**the information of shRNA used in this experiment.**

| shRNA name | sequence |
|---|---|
| shKDM3A | gatccGAGTGTGTGTGGATTGCTATCAAGAGTAGCAATCCACACACACTCTTTTTTg |
| shKDM3B | gatccCCTGTAACTTGACTGATACCTCAAGAGGGTATCAGTCAAGTTACAGGTTTTTTg |

**Supplementary table 3**

**the information of primers used in this experiment.**

| primer name | forward sequence | reverse sequence |
|---|---|---|
| ex-OSKM | TCGGACCACCTTGCCTTACAC | CAACGCCCAAAGGAAATCCAG |
| total-OCT4 | GTTTCAAGAACATGTGTAAGCTGCG | GATACTTGTCCGCTTTCTCTTCCG |
| total-SOX2 | GCAACCAGAAGAACAGCCCAGA | GTTGTGCATCTTGGGGTTCTCTTG |
| endo-OCT4 | CTTCACCACCCTGTACTCCTCG | CAGGCTTCTCTCCCTAGCTCAC |
| endo-SOX2 | ATGTCCCAGCACTACCAGAGCG | CTTACTCTCCTCCCATTTCCCTCT |
| endo-KLF4 | TGGGGGAGGGAAGACCAGAAT | TAGAACCAAGACTCACCAAGCACC |
| endo-c-MYC | GCAAAAGCTCGTGTCAGAAAAAG | TAGTTCCTCCCTCCAATAGGTCAAT |
| LIN28A | GAAGTCTGCTAAGGGCTTGGAATC | TGTCTCCCTTGGATCTGCGTTT |
| NANOG | CTTCACCAATGCCTGAGGTTTATG | AGGGCTGTCCTGAATAAGCAGATC |
| TAF4B | CCAGAAATGGGGCAGAATGTG | ACCAGGTGAGGCTGAGGTGAAG |
| SALL4 | AGAACAGCCGCACTGAGATGG | GCCGCTAACAACGGTGTCATAC |
| EF1-OCT4 | CTTCACCACCCTGTACTCCTCG | CGTGGGCTTGTACTCGGTCAT |
| EF1-SOX2 | ATGTCCCAGCACTACCAGAGCG | CGTGGGCTTGTACTCGGTCAT |
| EF1-KLF4 | ACACGAAGAGTTCTCATCTCAAGGC | CGTGGGCTTGTACTCGGTCAT |
| EF1-MYC | GCAAAAGCTCGTGTCAGAAAAAG | CGTGGGCTTGTACTCGGTCAT |
| EF1-LIN28A | CACAGGGAAAGCCAGCCTAC | CGTGGGCTTGTACTCGGTCAT |
| KDM3A | AAACACTGCTTCTGGCTCACTC | CTTTCAGCATAGCAACCGCATC |
| KDM3B | GGAAGCCAGAAGCCTTTAGCC | CCATCCCAGAAATCCCGAACT |
| KDM3C | TCCAAATAGCGGAACATCACCTC | TCCCATCGAATAAACTGGTGCA |
| KDM4A | CGCAAGATAGCCAATAGCGATAAG | CGTTCACATCCGCACCGTAGA |
| KDM4B | CCTGCGGTGGATTGATTACGG | GTGAGGTCTTTGCCCTGCTTC |
| KDM4C | GATGGCAAACTCTACGGGGC | CGCTTTCACTCTCTTGGGTAACTC |
| LIN28A-ChIP | AGCAGGACTGAAAGAAGGAAGGG | ATCAAGCAGGATTGGGCACC |
| OCT4-ChIP | GACAGACAAACATCATCCCCAGC | GGGTCTATTTTTCTGGGTCTGTCC |
| SALL4-ChIP | ATACAGTTGCTTGTGGAAAGTGCTT | GTTAAACTTCCAGTAAGGCTATGTCTC |
| TAF4B-ChIP-1 | CGGAAATCAGTTCTCCACATCG | GCGTTCTTCTTTCTCCTTGGTATCC |
| TAF4B-ChIP-2 | AGTTGGCTGTAGGTGTGTGGG | GCAGCAATCATCAGAACCGTAT |

**Supplementary table 4**

**the information of antibodies used in this experiment.**

| antibody | source | Catalog number |
|---|---|---|
| Histone H3K9me1 antibody (pAb) | Active motif | 39249 |
| Histone H3K9me2 antibody (pAb) | Active motif | 39375 |
| Histone H3K9me3 antibody (pAb) | Active motif | 39161 |
| KDM3A, JMJD1A Antibody | proteintech | 12835-1-AP |
| β-Actin mouse monoclonal antibody | sungene biotech | KM9001T |
| KDM3B Antibody | proteintech | 19915-1-AP |
| AFP Antibody | proteintech | 14550-1-AP |
| Desmin Antibody | proteintech | 16520-1-AP |
| TUBB3-Specific Antibody | proteintech | 66375-1-Ig |
| Human/Mouse Oct-3/4 Antibody | R&D Systems | AF1759 |
| Human/Mouse/Rat SOX2 Antibody | R&D Systems | AF2018 |
| Histone H3K4me1 antibody (pAb) | active motif | 39297 |
| Histone H3K4me2 antibody (pAb) | active motif | 39913 |
| Histone H3K4me3 antibody (pAb) | active motif | 39915 |
| Lin28a(C-9） | santa | sc-374460 |
| Monoclonal ANTI-FLAG® M2 antibody | sigma | F1804 |
| EZview™ Red ANTI-FLAG® M2 Affinity Gel | sigma | F2426 |
| Mouse Monoclonal Antibody to Human Histone H3 | Sino Biological | 100005-MM01 |
